# Prolonged epigenetic and synaptic plasticity alterations following single exposure to a psychedelic in mice

**DOI:** 10.1101/2021.02.24.432725

**Authors:** Mario de la Fuente Revenga, Bohan Zhu, Christopher A. Guevara, Lynette B. Naler, Justin M. Saunders, Zirui Zhou, Rudy Toneatti, Salvador Sierra, Jennifer T. Wolstenholme, Patrick M. Beardsley, George W. Huntley, Chang Lu, Javier González-Maeso

## Abstract

Clinical evidence suggests a potential therapeutic effect of classic psychedelics for the treatment of depression. The most outstanding and distinct characteristic is the rapid and sustained antidepressant action with one single exposure to the drug. However, the biological substrates and key mediators of psychedelics’ enduring action remain unknown. Here, we show that a single administration of the psychedelic DOI produced fast-acting effects on frontal cortex dendritic spine structure and acceleration of fear extinction via the 5-HT_2A_ receptor. Additionally, a single dose of DOI led to changes in chromatin organization particularly at enhancer regions of genes involved in synaptic assembly that stretched for days after the psychedelic exposure. DOI-induced alterations in neuronal epigenome overlapped with genetic loci associated with schizophrenia, depression and attention deficit hyperactivity disorder. Together, these data support the notion that epigenetic-driven changes in synaptic plasticity operate as the mechanistic substrate of psychedelic’s long-lasting antidepressant action but also warn on the limitations in individuals with underlying risk for psychosis.

## INTRODUCTION

Psychiatric conditions including depression, anxiety and stressor-related disorders affect the life of millions of individuals worldwide (*1, 2*). All currently available medications, including monoamine reuptake-based antidepressants such as fluoxetine, paroxetine and citalopram, require several weeks to months for the clinically relevant improvements to occur, and there is a high proportion of patients taking standard pharmacotherapies who remain treatment resistant (*3*). In addition, these pharmacological interventions are often accompanied by undesirable side effects (*4*). There is therefore an urgent need for better antidepressant treatments, with a faster onset of action, which will also offer relief to patients who do not respond to classical antidepressants.

Psychedelics, such as psilocybin, lysergic acid diethylamide (LSD) and the substituted amphetamine 1-(2,5-dimethoxy-4-iodophenyl)-2-aminopropane (DOI), are psychoactive compounds that profoundly affect various mental domains, particularly sensory perception and thought processes (*5*). Most of the previous preclinical and human neuroimaging studies have focused their efforts on the effects that occur within minutes to hours after psychedelic administration, with the aim of elucidating the molecular and neural circuit mechanisms responsible for their psychotic-like states (*6, 7*). However, recent pilot clinical trials suggest that psychedelics may represent a promising long-lasting treatment for patients with depression and other psychiatric conditions (*8–11*). As an example, cancer patients demonstrated rapid increases in positive affect after psilocybin administration, with 60-85% showing long-term improvements in anxiety and depression measures at a 6.5-month follow-up (*11*). Despite these striking effects, their acute psychotic symptoms and drug abuse potential preclude the routine use of psilocybin and other psychedelics in daily clinical practice. Therefore, there is a clear need for basic and translational research focused on understanding the molecular mechanisms mediating the clinical effectiveness of psychedelics – with the ultimate goal of developing safer, more effective, and non-psychedelic treatment strategies.

Preclinical assays in rodent models have evaluated the effects of psychedelics as rapid-acting antidepressant medications. Notably, it has been suggested that a single administration of psilocybin or LSD produces long-lasting (for 5 weeks post-administration) antidepressant-like effects in rats within models of behavioral despair or passivity such as the forced swim test (*12*). Additionally, it has been reported that a single dose of the psychedelic *N,N*-dimethyltryptamine (DMT) facilitates the extinction of cued fear memory in rats (*13*). The pharmacological profiles of these drugs are notoriously complex (*14*), but it is well established that the serotonin 5-HT_2A_ receptor (5-HT_2A_R) in the frontal cortex is involved in the effects of psychedelics on psychosis-like behavior (*15*). With regard to outcomes relevant to depression, anxiety and stressor-related disorders, the receptor targets responsible for these benign effects of psychedelics remain largely unexplored.

Here, we aimed to elucidate the molecular mechanism responsible for the post-acute effects of the psychedelic phenethylamine DOI using mouse behavior models relevant to depression, anxiety and stressor-related disorders, as well as its implication in crucial plasticity phenotypes including frontal cortex dendritic spine structure and gene expression.

## RESULTS

### Psychedelics accelerate fear extinction via 5-HT_2A_R

To explore the potential role of 5-HT_2A_R in the post-acute effects of psychedelics on behavior models relevant to anxiety, passivity, cognition and sensorimotor gating, mice were tested 24 h after administration of the psychedelic yet relatively selective 5-HT_2A/2C_R agonist DOI (*16*), or vehicle – a timepoint at which DOI is no longer detectable in the mouse brain (*17*). Locomotor activity in a novel environment as a model of exploratory behavior (Figs. 1A and 1B) and time spent in the light compartment using a dark-light choice test as a conflict paradigm (Figs. 1C and 1D) were similar between DOI-treated and vehicle-treated mice. Novel-object recognition (Fig. 1E) and prepulse inhibition of startle (Fig. 1F) were also unaffected by DOI treatment. However, immobility time in the forced swim test was reduced in DOI-treated mice as compared to controls (Fig. 1G). These findings indicate that a single dose of DOI reduces behavioral despair or passivity as a model of depression 24 h after its administration, whereas behaviors in models of anxiety, recognition memory and sensorimotor gating remain unchanged.

**Fig. 1.**
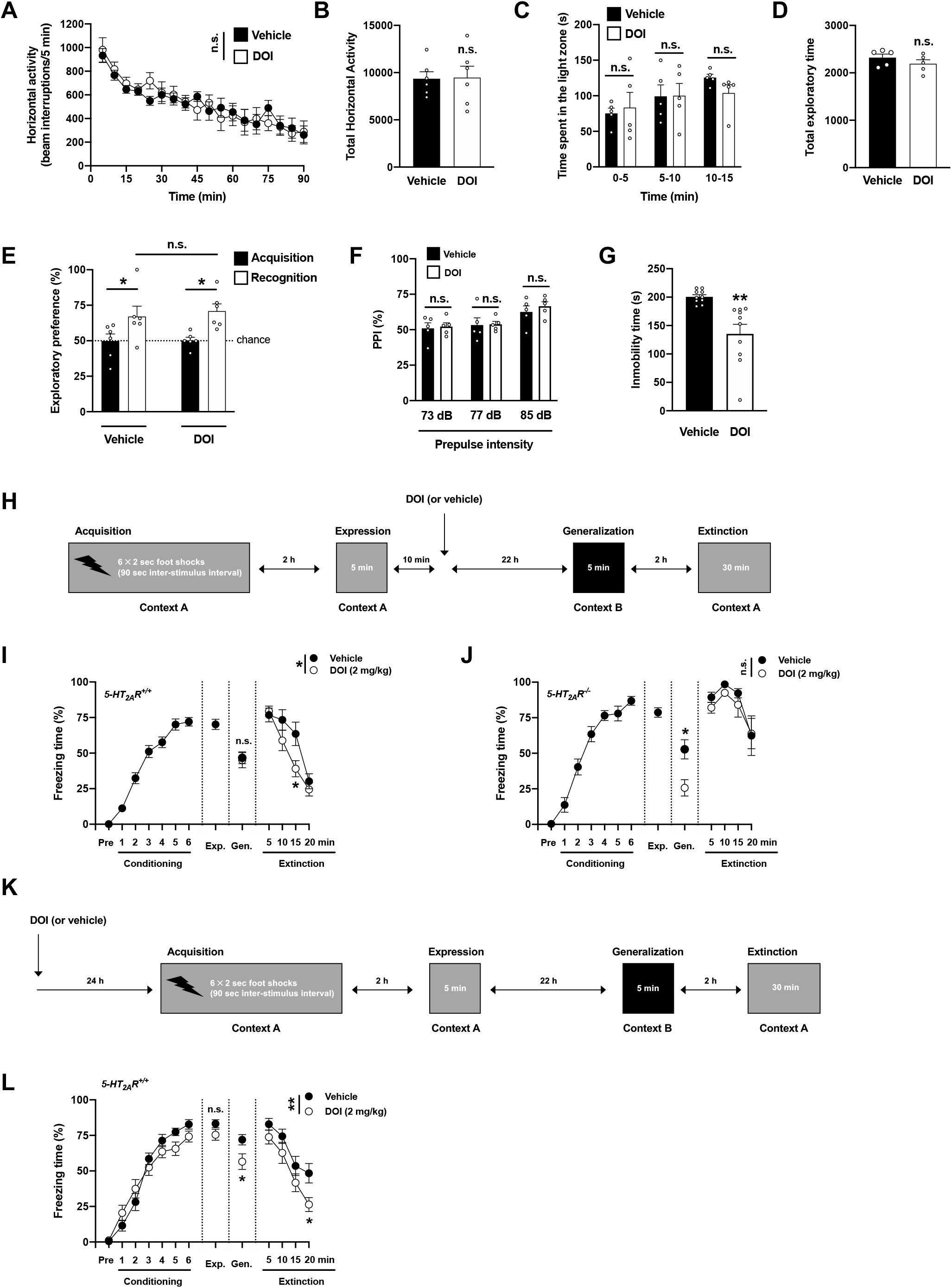
Post-acute effects of DOI on passivity and fear extinction. **(A-G)** Behavior was tested 24-h after a single injection (i.p.) of DOI (2 mg/kg) or vehicle. **(A,B)** Lack of effect of DOI on exploratory behavior in an open field (n = 6 mice per group). Time-course (**A,** F[1,10] = 0.006, p > 0.05), and total horizontal activity (**B,***t*_10_ = 0.07, p > 0.05). (**C,D**) Lack of effect of DOI on dark-light choice test (n = 5 mice per group). Time (s) spent in the light zone (**C,** F[1,8] = 0.08, p > 0.05), and total exploratory time during the 15-min test (**D,***t*_8_ = 1.14, p > 0.05). **(E)** Lack of effect of DOI on the novel-object recognition test (n = 6 mice per group, acquisition vs recognition, F[1,20] = 13.06, p < 0.01; vehicle vs DOI, F[1,20] = 0.16, p > 0.05). **(F)** Lack of effect of DOI on the PPI of startle (n = 5 mice per group, F[1,24] = 0.41, p > 0.05). **(G)** Reduction of immobility time (s) during the last 4 min of the 6-min forced swimming test (10 mice per group, *t*_18_ = 3.80; p < 0.01). (**H-J**) Effect of DOI on contextual fear extinction in *5-HT_2A_R^+/+^* (**I**) and *5-HT_2A_R^−/−^* (**J**) mice. Timeline of the experimental design (**H**). Fear conditioning (**I,** n = 26 mice, F[6, 175] = 73.69, p < 0.001; **J,** n = 12 mice, F[6, 77] = 58.76, p < 0.001), generalization (**I,** n = 11-15 mice per group, *t*_24_ = 0.24, p > 0.05; **J,** n = 6 mice per group, *t*_10_ = 3.04, p < 0.05), extinction (**I,** n = 11-15 mice per group, F[1, 96] = 5.51, p < 0.05; **J,** n = 6 mice per group, F[1, 40] = 2.29, p > 0.05). (**K,L**) Effect of DOI on fear acquisition. Timeline of the experimental design (**K**). Fear conditioning (n = 14-15 mice per group; conditioning, F[6,189] = 95.86, p < 0.001; vehicle vs DOI, F[1,189] = 0.91, p > 0.05), expression (n = 14-15 mice per group, *t*_27_ = 1.61, p > 0.05), generalization (n = 14-15 mice per group, *t*_27_ = 2.34, p < 0.05), extinction (n = 14-15 mice per group; extinction, F[3,104] = 20.17, p < 0.001; vehicle vs DOI, F[1,104] = 10.65, p < 0.01) (**L**). Statistical analysis was performed using two-way repeated measures ANOVA (**a**), two-way ANOVA with Sidak’s multiple comparison test (**C, D, F, I, J, L**), or Student’s *t* test (**B, D, G, I, J, L**). *p < 0.05, **p < 0.01, ***p < 0.001, n.s., not significant.

Fear and anxiety are adaptive defensive behaviors that are mediated by different neural substrates (*18*). To test whether DOI affects contextual fear conditioning and extinction, as well as the role of 5-HT_2A_R-dependent signaling in these effects, *5-HT_2A_R^+/+^* and *5-HT_2A_R^−/−^* mice were tested in a paradigm of fear acquisition (context A) and expression (context A). After this, mice were randomly separated into two groups that received a single dose of DOI or vehicle. The day after, DOI- and vehicle-treated groups were tested for fear generalization (context B) and context fear extinction (context A) (Fig. 1H). Our data show that there is a modest yet statistically significant increase in fear during fear acquisition in *5-HT_2A_R^−/−^* mice as compared to *5-HT_2A_R*^+/+^ controls (fig. S1), whereas no differences were observed during the fear expression session (fig. S1). Administration of DOI had no effect on fear generalization in *5-HT_2A_R^+/+^* mice, but reduced freezing time during the generalization session in *5-HT_2A_R^−/−^* littermates (Figs. 1I and 1J). Fear extinction was significantly reduced in vehicle-treated *5-HT_2A_R^−/−^* mice as compared to vehicle-treated *5-HT_2A_R*^+/+^ controls (fig. S1). Additionally, *5-HT_2A_R^+/+^* mice that had previously received a single dose of DOI showed faster development of contextual fear extinction as exemplified by the progressive reduction in freezing time compared to vehicle-treated animals (Figs. 1I and 1J) – an effect that was not observed in *5-HT_2A_R^−/−^* littermates (Figs. 1I and 1J). To test whether psychedelics affect fear acquisition, a different cohort of *5-HT_2A_R^+/+^* mice was subjected to a similar paradigm of fear acquisition, expression, generalization and extinction but with DOI or vehicle being administered 24 h before the fear acquisition protocol (Fig. 1K). Our data show that DOI did not affect fear acquisition or fear expression, whereas it reduced freezing time during the fear generalization and context fear extinction sessions (Fig. 1L) – suggesting that the effects of DOI on associative learning relevant to contextual fear extinction may linger beyond the 24h after drug administration.

### Psychedelics enhance cortical dendritic density via 5-HT_2A_R

Structural and functional modification of dendritic spines are central to brain plasticity (*19, 20*). As a general rule, increased synaptogenesis and functional plasticity are tightly correlated with the size and the shape of a dendritic spine; stubby spines are theorized to be transitional structures that will enlarge, possibly into mature mushroom spines, whereas thin spines are highly dynamic and likely to change in response to activity. Basic and clinical studies demonstrate that depression is associated with reduction of synaptic density in brain regions that regulate anxiety, mood and fear extinction, including the frontal cortex (*3, 21*). Typical antidepressants like fluoxetine can reverse these synaptic deficits, albeit with limited efficacy and delayed response. Some (*22*) but not all (*23, 24*) previous studies suggest that psychedelics promote rapid structural plasticity in pyramidal neurons. Additionally, most of these later data were obtained in vitro in neuronal primary cultures or relied upon manual 2D images after Golgi staining. This is important because previous studies have demonstrated dramatic density differences between 2D Golgi and 3D methods and also a selective bias against immature thin spines in 2D counts (*25, 26*). Here we measured the effect of a single administration of DOI or vehicle on dendritic spine density in cortical pyramidal neurons in *5-HT_2A_R^−/−^* mice and controls. We targeted pyramidal neurons from frontal cortex of adult mice through the injection of adeno-associated virus (AAV8) encoding enhanced yellow fluorescent protein (eYFP) under the control of the *CaMKIIα* promoter. Using a 3D automated method for quantitative structural spine analysis, our findings suggest a lower spine density in the frontal cortex of vehicle-treated *5-HT_2A_R^−/−^* mice as compared to *5-HT_2A_R^+/+^* controls (Figs. 2A and 2B), a phenotype that was primarily driven by a selective decrease of stubby (Figs. 2A and 2C) and thin (Figs. 2A and 2D), but not mushroom (Figs. 2A and 2E), spines. Additionally, a single administration of DOI selectively augmented the density of transitional stubby (Figs. 2A and 2C) and dynamic thin (Figs. 2A and 2D) spines in CaMKIIα-positive frontal cortex neurons, but not mature mushroom spine density (Figs. 2A and 2E). This psychedelic-induced synaptic remodeling event required expression of 5-HT_2A_R, as the effect of DOI on dendritic spine structure was not observed in *5-HT_2A_R^−/−^* mice (Figs. 2A-2E).

**Fig. 2.**
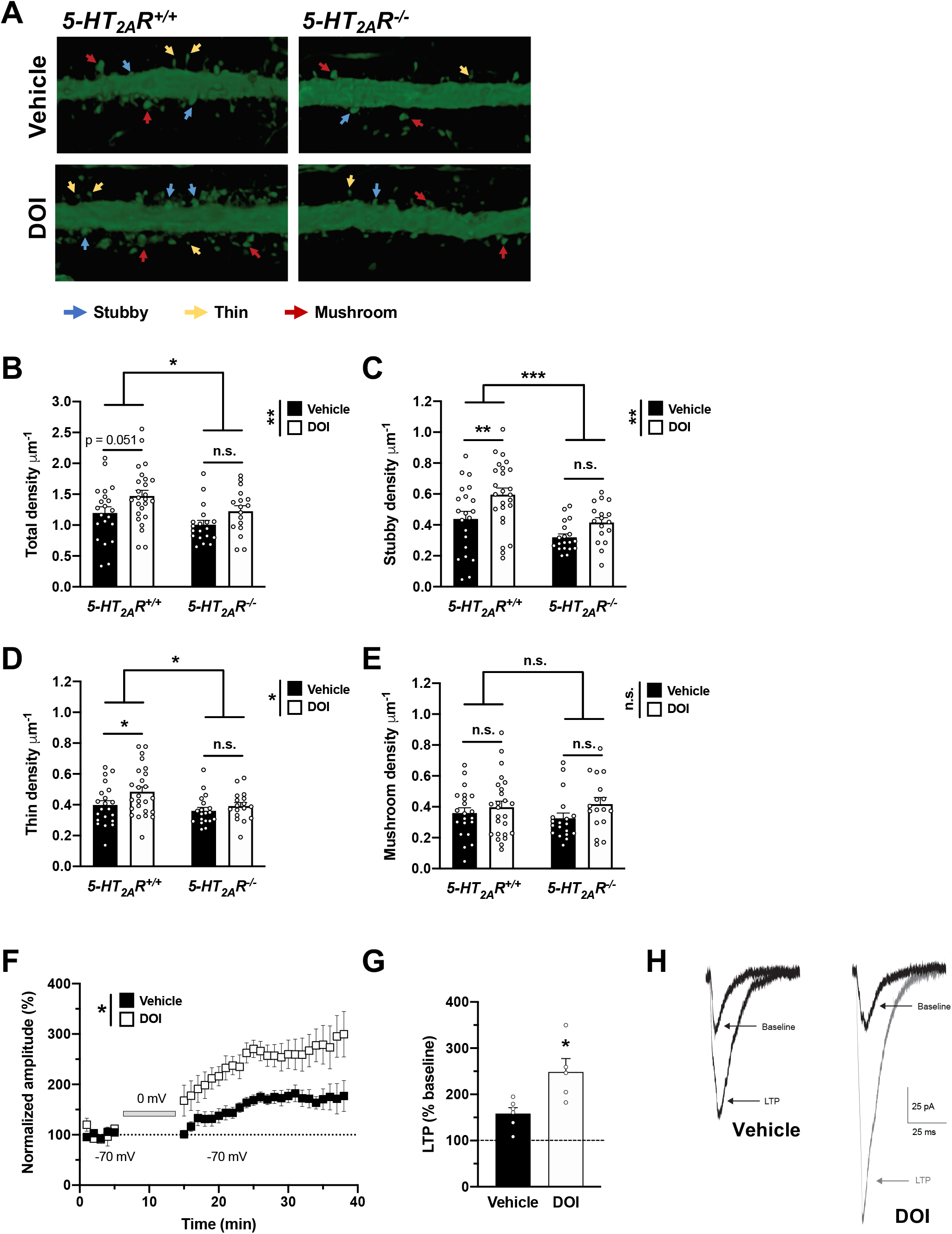
Post-acute effects of DOI on frontal cortex synaptic plasticity. (**A-D**) Effect of DOI on synaptic structural elements in the frontal cortex of *5-HT_2A_R^+/+^* and *5-HT_2A_R^−/−^* mice (n = 17-25 dendrites from independent neurons from both hemispheres in 3-4 mice per group). Samples were collected 24-h after a single injection (i.p.) of DOI (2 mg/kg) or vehicle. Representative three-dimensional reconstructions of AAV-injected cortical dendritic segments (**A**). Total (**B**, *5-HT_2A_R^+/+^* vs *5-HT_2A_R^−/−^* mice, F[1,78] = 5.72, p < 0.05; vehicle vs DOI, F[1,78] = 7.31, p < 0.01), stubby (**C**, *5-HT_2A_R^+/+^* vs *5-HT_2A_R^−/−^* mice, F[1,78] = 13.56, p < 0.001; vehicle vs DOI, F[1,78] = 9.65, p < 0.01), thin (**D**, *5-HT_2A_R^+/+^* vs *5-HT_2A_R^−/−^* mice, F[1,78] = 5.27, p < 0.05; vehicle vs DOI, F[1,78] = 4.28, p < 0.05), mushroom (**E**, *5-HT_2A_R^+/+^* vs *5-HT_2A_R^−/−^* mice, F[1,78] = 0.03, p > 0.05; vehicle vs DOI, F[1,78] =2.90, p > 0.05). (**F-H**) A single dose (i.p) of DOI (2 mg/kg) significantly enhanced cortical LTP in comparison with saline-injected mice assayed 24-h post-injection. Normalized EPSC amplitudes obtained from whole-cell patch-clamp recordings of L2/3 neurons from either DOI-injected (n = 5 neurons from 5 mice) or vehicle-injected (n = 6 neurons from 5 mice) animals. LTP was induced by a protocol that pairs extracellular presynaptic stimulation of L4 neurons with brief (10 min) postsynaptic depolarization of overlying L2/3 neurons to 0 mV (gray bar). Symbols represent EPSCs averaged by 1 min bins (vehicle vs DOI, F[1,10] = 9.84, p < 0.05) (**F**). Average magnitude of LTP (normalized EPSCs recorded over the 15-40 min post-induction period) from DOI-treated mice and controls (*t*_9_ = 3.05, p < 0.05) (**G**). Representative traces of EPSCs recorded from L2/3 neurons from DOI- or vehicle-treated mice at baseline and following induction of LTP (**H**). Statistical analysis was performed using two-way ANOVA with Sidak’s multiple comparison test (**B, C, D, E**), two-way repeated measures ANOVA (**F**), or Student’s *t* test (**G**). *p < 0.05, **p < 0.01, ***p < 0.001, n.s., not significant.

Based on these findings of dendritic spine structural plasticity, we next determined whether DOI also regulates persistent synaptic functional plasticity in frontal cortex pyramidal neurons. In vehicle-treated mice, as expected, pairing layer 4 (L4) stimulation with brief postsynaptic depolarization induced robust long-term potentiation (LTP) in L2/3 neurons that was sustained for 30-40 min until the experiment was terminated (Figs. 2F-2H). Notably, a significantly greater magnitude of LTP was evident in DOI-treated mice in comparison with that in vehicle-treated mice over the same time-course (Figs. 2F-2H), indicating that DOI enhanced synaptic plasticity in L2/3 frontal cortex pyramidal neurons.

### Psychedelics lead to long-lasting alterations in frontal cortex epigenomic landscapes

Using classical gene expression assays such as microarrays, previous studies demonstrated that a single administration of psychedelics including DOI and LSD alters the level of expression of several genes that showed maximal changes at ~60 min – returning to basal level at ~2 h after drug exposure (*27, 28*). Several of the genes showing transient induction upon acute psychedelic administration have previously been implicated in processes related to transcription and chromatin organization. However, these previous transcriptomic studies in bulk tissue samples suffered from lack of cell type specificity in their profiling. Here we tested whether a single dose of DOI leads to long-lasting epigenomic and transcriptomic alterations in the frontal cortex using high-resolution, cell type-specific and low-input ChIP-seq and RNA-seq measurements. Mice were injected with DOI, or vehicle, and frontal cortex samples were collected 24h, 48h, or 7d after drug (or vehicle) administration (fig. S2). Histone modification H3K27ac (acetylation of histone H3 at lysine 27) along with the transcriptome were profiled in NeuN-positive (NeuN^+^) neuronal nuclei isolated from the frontal cortex by fluorescence-activated cell sorting (FACS). We applied <u>M</u>icrofluidic <u>O</u>scillatory <u>W</u>ashing ChIP-seq (MOWChIP-seq) (*29, 30*) and Smart-seq2 (*31, 32*) to profile H3K27ac and transcriptome, respectively, using the small quantity of neuronal nuclei yielded from the mouse frontal cortex (table S1).

Enhancers are highly dynamic epigenomic regulatory elements with known involvement in plasticity and neurodevelopmental processes (*33*). We predicted enhancers by scanning the H3K27ac^high^ regions that did not intersect with promoters. We conducted K-means clustering of dynamic enhancers, and examined gene ontology (GO) terms and enriched transcription factor (TF) binding motifs associated with the clusters (Fig. 3A). We divided all dynamic enhancers into 6 clusters with various patterns of variation after DOI injection (Fig. 3A). Clusters 1-3 represent enhancers exhibiting eventual increase after DOI administration, while clusters 4-6 show a decrease in DOI-injected mice as compared to vehicle-treated mice. Clusters 1-2 show an increase that peaks at 48 h before decreasing at 7 d, while cluster 3 experiences a much slower increase that shows the highest intensity at 7 d. Clusters 4-5 have the deepest decrease at 24h while enhancer intensity slowly recovers over 48 h and 7d. Cluster 6 presents an initial decrease at 24 h followed by minor fluctuations at 48 h and 7 d. Interestingly, at least 32.7% of these dynamic enhancers (clusters 3 and 6) are still in their altered state at 7 d, suggesting long-lasting effects of DOI at the epigenomic level that outlast by several days the presence of the drug in native tissue.

**Fig. 3.**
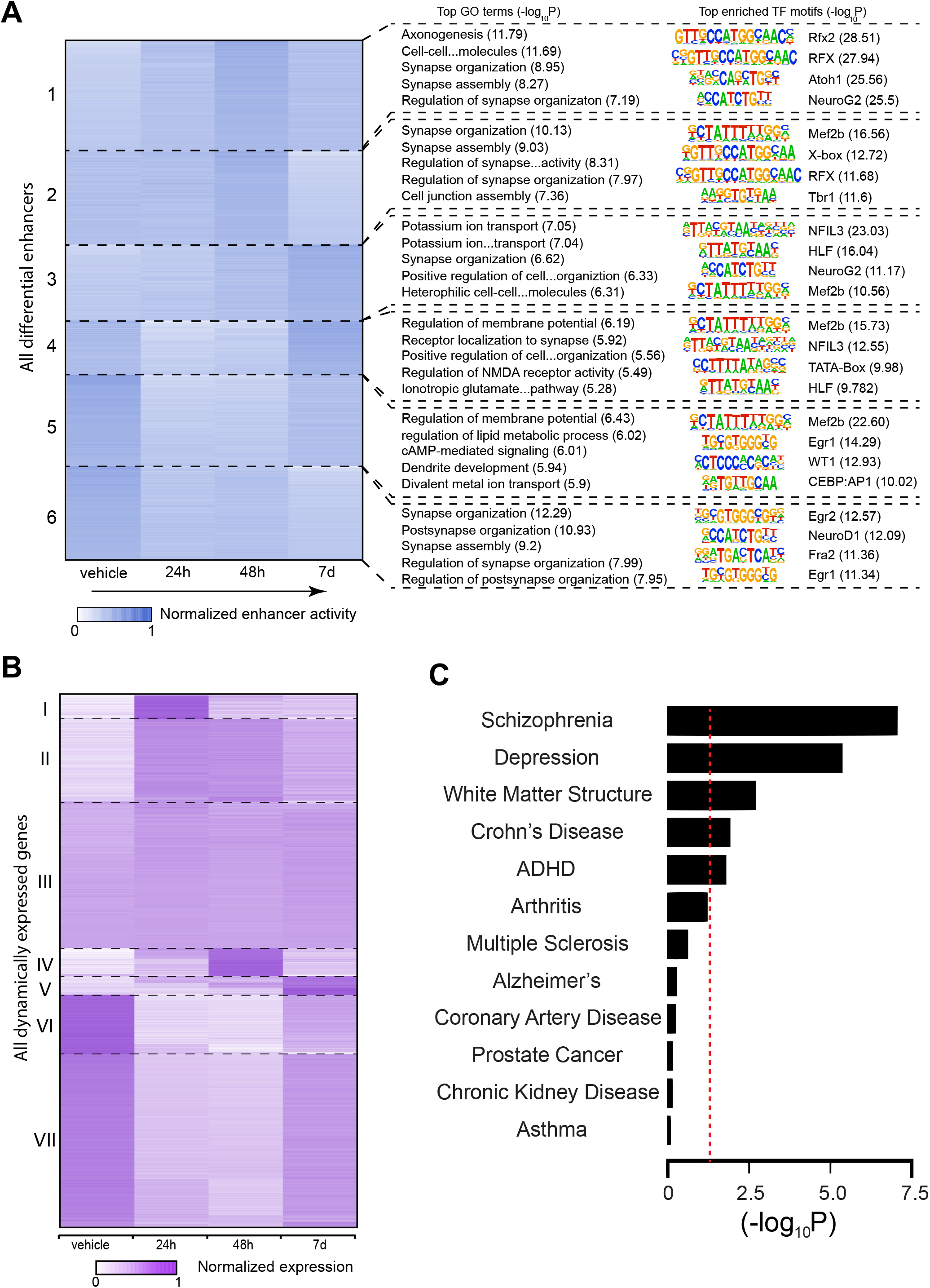
Post-acute effects of DOI on frontal cortex epigenomic and transcriptomic variations. **(A)** Effect of DOI on time-lapse epigenomic and transcriptomic variations in neuronal nuclei of the mouse frontal cortex. K-means clustering of differential enhancers based on normalized H3K27ac signal (n = 3995; Cluster sizes from top to bottom: 852, 718, 585, 419, 700, 721). The top five enriched biological process gene ontology (GO) terms identified by clusterprofiler are listed next to each cluster, and the top four motifs enriched in each cluster are listed next to the GO terms together with their sequences **(B)** K-means clustering of normalized gene expression (n = 13605). Cluster sizes from top to bottom: 604, 2120, 3748, 708, 497, 1507, 4421. **(C)** Significance of overlap between differential H3K27ac peaks and NHGRI-EBI GWAS SNP sets. Significance is calculated using a null distribution and is shown as -log(p-value). The red dotted line denotes p < 0.05 cutoff. GWAS sets are ordered according to significance. Frontal cortex samples were collected 24-h, 48-h, and 7-d after a single injection (i.p.) of DOI (2 mg/kg) or vehicle.

To directly interrogate the functional relevance of these findings, we examined the GO terms and TF motifs associated with each of the clusters (Fig. 3B and tables S2-S3). Synapse organization and assembly are enriched in all clusters to various degrees (table S2), consistent with the effects of DOI on structural and synaptic plasticity. The top GO terms enriched in clusters 1 and 2 include axonogenesis, synapse organization, and cell junction assembly. Motifs of several TFs importantly involved in neuronal growth and synaptic plasticity, such as *Mef2B*, *NeuroG2* and *Atoh1* (*34*), are also highly enriched in these clusters. Clusters 4 to 6, which experience a decrease upon DOI administration, are associated with GO terms related to regulation of glutamate NMDA receptor activity, regulation of membrane potential, and synapse assembly. Enrichment of two zinc finger-containing transcription factors, *Egr1* and *Egr2*, also occurs in clusters 5 and 6. Considering previous findings showing that psychedelics lead to an acute increase in glutamate release as well as augmentation of *Egr1* and *Egr2* transcription in mouse frontal cortex neurons (*15*), these data suggest a compensatory mechanism as a potential explanation for the enhancer intensity decrease observed in clusters 4 to 6 upon DOI administration. Clusters 3 and 6 represent enhancers with long-lasting (7d) DOI-dependent increased and decreased activity, respectively. Cluster 3 shows enrichment in GO terms associated with rhythmic processes. This finding, together with *NFIL3*, a TF involved in the mammalian circadian oscillatory mechanism (*35*), as the top enriched TF motif in this cluster, shows agreement with long-term effects of psychedelics on sleep-wake cycles. Cluster 6 has synaptic plasticity-related genes heavily enriched (table S2).

The transcriptomic variations induced by DOI appear to be much more transient than the epigenomic changes (Fig. 3B). Based on K-means clustering, only 3.7% (cluster V) of the dynamically expressed genes (DEGs) present long-term variation (7 d), and only a small number of these genes (n = 574, or 4.2% of the total DEGs) exhibit a pattern of expression that correlates with enhancer dynamics (*i.e*., with a Spearman’s correlation r > 0.4 between enhancer intensity and expression level) (fig. S3 and table S4). These results suggest that the long-term effects of the psychedelic DOI are associated with epigenomic regulations of greater magnitude than changes in transcriptomic dynamics.

To determine if there was a significant overlap of differential H3K27ac peaks with single nucleotide polymorphisms (SNPs) associated with depression that were reported based on genome wide association studies (GWAS), a null distribution was calculated using the set of all SNPs (Fig. 3C). This was repeated for different GWAS sets that are associated with various traits to determine specificity. Of twelve GWAS loci datasets, five (schizophrenia, depression, white matter structure, Crohn’s disease and attention deficit hyperactivity disorder [ADHD]) were statistically significant (Fig. 3C and table S5).

We further investigated our RNA-seq data using weighted correlation network analysis (WGCNA) with the goal of detecting co-expression modules with highly correlated genes to associate these modules with sample traits or experimental groups (*36*). 14,727 genes with their expression levels throughout all samples in the top 30% were selected and the generated data matrix was used as the input for WGCNA. After construction of a co-expression network from the matrix, ten modules with densely interconnected genes were detected and the expression pattern in each module was summarized by the module eigengene (table S5). The eigengene significance was used for assessing correlation between experimental groups and each module. We performed GO analysis on the genes included in these modules and 5 modules (blue, green, turquoise, yellow, and magenta) showed significantly enriched GO terms (Figs. 4B, 4E and figs. S4B, S4E, S4H).

**Fig. 4.**
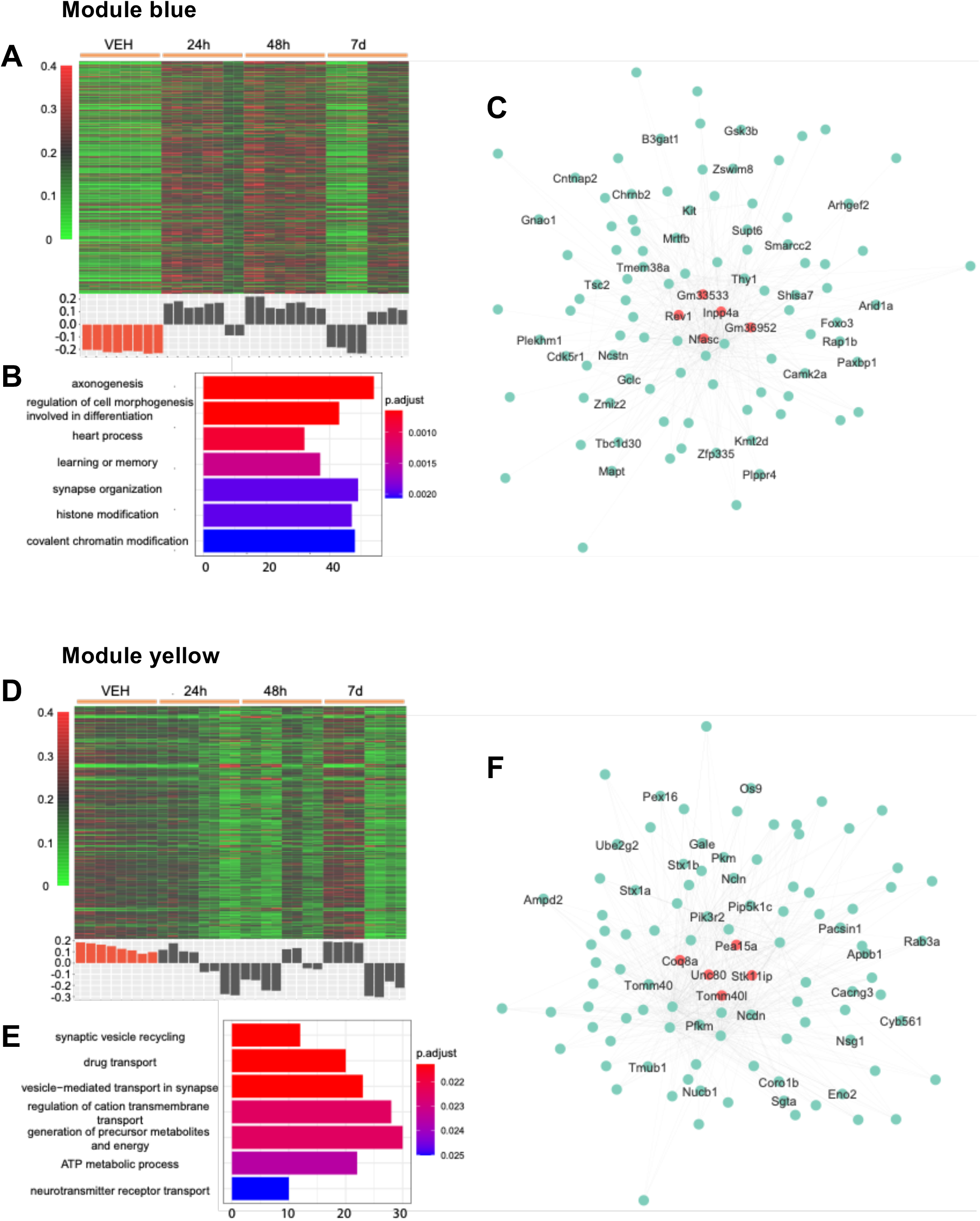
Two gene co-expression modules (blue and yellow) associated with administration of DOI. **(A,D)** Heatmap of normalized gene expression profiles in the co-expression module (top). The module eigengene values across samples in four experimental groups (bottom). **(B,E)** Selected top categories from GO enrichment analysis. **(C,F)** Visualization of the intramodular connections among the top 100 hub genes in each module. The top 5 genes are in large size and colored orange. The genes involved in the top 15 GO terms are labeled.

Among all 10 modules, module blue showed the strongest positive correlation with group 48 h and the strongest negative correlation with the control group. Compared to the control, the eigengene of module blue is overexpressed in both 24h and 48h groups before decreasing to the control level on 7d (fig. S4J). GO analysis of this cluster of 1933 genes leads to terms including: “*Axonogenesis*” (p.adjust 6.44 × 10^−4^), “*Heart process*” (p.adjust 8.32 × 10^−4^), “*Learning or memory*” (p.adjust 1.35 × 10^−3^), and “*Histone modification*” (p.adjust 1.95 × × 10^−3^) (Fig. 4B). The GO term “*Axonogenesis*” is also identified in the cluster 1 of dynamic enhancers (Fig. 3A). *Inpp4a*, *Nfasc*, and *Cntnap2* are in the top 100 hub genes of module blue (table S6). Previous studies suggested that *Inpp4a* may play an important role in the development of epilepsy (*37*). Epileptic seizures have been reported as a result of psychostimulant drug use, although this adverse effect is less common with psychedelics than with other psychostimulants such as cocaine (*38*). Loss of function of *Inpp4a* also increases severity of asthma, which may explain the mechanism underlying the effect of DOI preventing allergic asthma in a mouse model (*39*). *Cntnap2* and *Nfasc*, which are genes previously involved in axonal guidance, synaptogenesis and neuron-glial cell interactions (*40–42*) as well as implicated in neurodevelopmental psychiatric conditions such as schizophrenia and autism (*43–45*), were also identified in differential enhancers from clusters 3 and 6 (Fig. 3A), respectively.

The eigengene expression in module yellow shows opposite trend of that in module blue (Fig. 4D). Among all 10 modules, module yellow is the only one that showed a positive correlation with the control group (fig. S4J). GO biological process enrichment for the genes in module yellow leads to terms including: “*Synaptic vesicle recycling*” (p.adjust 0.021), “*Drug transport*” (p.adjust 0.021), “*Vesicle-mediated transport in synapse*” (p.adjust 0.021), and “*Regulation of cation transmembrane transport*” (p.adjust 0.022). Interestingly, this module was also enriched with genes related to hypoxia-inducible factor 1 (HIF-1) signaling pathway, including *Slc2a1*, *Eno2* (hub gene), *Gapdh*, *Aldoa*, *Mapk3*, *Pfkl*, and *Pfkm* (hub gene). Since the HIF-1 pathway is a major modulator of hypoxia stress response pathways (*46*), these findings further support the hypothesis the 5-HT_2A_R modulate the ventilatory response to hypoxia (*47*).

Module magenta’s eigengene shows no correlation with any experimental groups (correlation coefficient in the range of −0.12 to 0.05, fig. S4J). The genes in module magenta are enriched in terms related to inflammatory response and intrinsic apoptotic signaling pathways. Modules green and turquoise have their eigengenes in a similar pattern as that of module blue. Their top GO terms were “*Synapse organization*” (p.adjust 1.89 × 10^−4^) and “*mRNA processing*” (p.adjust 2.61 × × 10^−17^).

## DISCUSSION

Previous observations in rodent models clearly showed that a single administration of psychedelics leads to long-lasting effects on synaptic plasticity and behavior models of depression (*12, 13, 22*). The manifestation of these phenotypes assumed, but never clearly demonstrated, that 5-HT_2A_R is the main molecular target responsible for the effects of psychedelics. In the present study, our data provide the first direct evidence that the post-acute effects of the psychedelic DOI on mouse frontal cortex dendritic spine structure and contextual fear extinction is 5-HT_2A_R-dependent. Our data also suggest that a single administration of DOI leads to long-lasting alterations in frontal cortex gene expression and chromatin organization that outlast the acute action of this psychedelic and its presence in the organism.

One of the still open questions in the field is whether the subjective effects of psychedelics are necessary or complementary for their post-acute clinically relevant outcomes in patients with severe psychiatric disorders such as depression (*48, 49*). Thus, it could be speculated that the fast-acting and long-lasting antidepressant properties induced by psychedelics is a consequence of complex psychological processes, such as waking consciousness, derealization and diminished ego-functioning (*50, 51*). It has also been reported that psilocybin can induce mystical and spiritually significant experiences to which participants attribute increases in well-being (*52*). These subjective components of psychedelic drug action appear to be difficult to control with placebo in the clinical setting due to the overwhelming nature of the drug effect. Accordingly, recent studies highlight the role for positive expectancy in predicting positive outcomes following psychedelic microdosing in healthy volunteers and increase suggestibility under the effect of these drugs (*53*). Obviously, these intricate mental experiences, which rely upon subjective reports, would be difficult, perhaps impossible, to model in rodents. However, the recursiveness of the neocortex structure across the mammalian clade and some evolutionarily preserved behaviors inherently linked to constructs relevant to mental health offer an unparalleled platform for characterizing non-subjective effects of psychedelics. Our data in this study, along with previous observations (*12, 13, 22*), suggest that favorable outcomes of psychedelics in processes related to cortical dendritic spine structure and behavior models relevant to depression and stressor-related disorders occur independently of subjective effects. It is then feasible that quantifiable molecular events and subjective experience concur in the therapeutic benefit of psychedelics. Studies like the present utilizing animal models can therefore offer valuable insights on the biological substrate of psychedelics’ therapeutic potential.

We found that a single administration of the psychedelic DOI leads to a post-acute (24 h after) increase on the density of immature and transitional dendritic spines that included thin and stubby, whereas the density of mature mushroom spines was unaffected. This effect of DOI on frontal cortex dendritic spine structure was not observed in *5-HT_2A_R^−/−^* mice, but it is important to remark that vehicle-injected *5-HT_2A_R^−/−^* mice also showed reduced density of frontal cortex thin and stubby dendritic spine density in the frontal cortex as compared to vehicle-injected *5-HT_2A_R^+/+^* controls. Although further investigation is necessary to unravel the mechanisms underlying this alteration in mice lacking the 5-HT_2A_R, a potential explanation may be related to changes among developmental compensatory pathways also involved in the alterations in fear conditioning and extinction observed in *5-HT_2A_R^−/−^* mice.

Our findings on the epigenomic and transcriptomic dynamics following DOI administration are significant. There have been no integrative studies of temporal changes in frontal cortex epigenome and transcriptome after psychedelic administration previously. Genome-wide epigenetic changes have long been speculated to be the layer of regulation that integrates both genetic and environmental factors (*54, 55*). In contrast with the determinism of genetic architecture, we show that the epigenetic landscape can be exogenously reshaped by a drug that elicits unequivocal psychedelic effects in humans. At the genomic level, genetic variations in non-coding regions may lead to propensity for depression, schizophrenia and other psychiatric conditions via long-range chromatin regulation such as enhancer-promoter interaction In this study, we report that, in contrast to the fairly rapid and transient changes in the transcriptome, a large fraction of epigenetic changes in enhancer regions persist for at least 7 d after DOI administration and potentially constitute the molecular basis for the long-lasting effects.

We also studied the points of convergence between genetic loci associated with a number of human disorders and the footprint of DOI in the mouse brain epigenome. The differential peaks of H3K27ac showed significant overlap with GWAS-discovered variants associated with schizophrenia, depression, and ADHD. Although psychedelics have been shown to have fast-acting and long-lasting therapeutic effect on depression (*21*), their effects on schizophrenia and ADHD remain unknown even when taking into account some overlap between these three psychiatric disorders (*56–58*). The acute effects of psychedelics resemble some of the positive symptoms (*i.e*., hallucinations and delusions) in patients with schizophrenia (*50, 51*), and chronic LSD administration alters rat frontal cortex expression of a group of genes relevant to schizophrenia (*59*). Markers linked to white matter structural changes, which have been associated with depression (*60*), were also significantly altered upon DOI administration. We identified genetic substrates associated with depression in the epigenomic landscapes of DOI exposure, but the concurrence of genetic markers for schizophrenia may serve as a molecular warning on the risks associated to the use of psychedelics in psychotic-prone individuals.

Other than outcomes in the treatment of depression, synaptic plasticity events in diverse parts of the brain have been linked to different adaptive and maladaptive traits. Previous studies have clearly demonstrated that psychoactive drugs of abuse such as cocaine increase dendritic spine density within key components of the brain’s reward circuitry, such as the nucleus accumbens (*61*). It has also been reported that the density of immature thin dendritic spines is increased in the frontal cortex of a genetic rat model that exhibits schizophrenia-relevant features (*62*). Brain regions enriched in 5-HT_2A_R include the frontal cortex, ventral striatum, several thalamic nuclei and the hypothalamus (*63, 64*). Considering our previous findings suggesting that the 5-HT_2A_R in frontal cortex pyramidal neurons is necessary and sufficient for the effects of psychedelics on psychosis-related behavior (*15*), here we focused our efforts on the post-acute effects of the psychedelic DOI in this particular brain region. Although our data provide the first evidence of long-lasting effects of a single administration of psychedelics on epigenomic landscape within neuronal nuclei in the frontal cortex, additional efforts will be required to establish whether these changes in synaptic plasticity and chromatin arrangement can be extrapolated to other brain regions relevant to substance abuse, and whether other cortical and subcortical dendritic structural and epigenetic plasticity events contribute to either clinically beneficial endpoints or, alternatively, unwanted side effects such as drug addiction, psychotic symptoms or hallucinogen persisting perception disorder.

Our previous work focused on the fundamental paradox that all known psychedelics are 5-HT_2A_R agonists, but 5-HT_2A_R agonism is not a sufficient condition for psychedelic action – as exemplified by structurally related drugs such as lisuride or ergotamine that do not exhibit psychedelic activity. We reported that a single dose of psychedelics, such DOI, LSD or psilocin, induces frontal cortex 5-HT_2A_R-dependent expression of genes associated with cell morphogenesis, neuron projection and synapse structure, such as *Egr1*, *Egr2*, *I_κ_B_α_*, and *N10*. Other genes, such as *c-Fos*, were induced by either psychedelic or non-psychedelic 5-HT_2A_R agonists (*15, 27*). Our data here support a scheme whereby a single dose of the phenethylamine DOI leads to changes in frontal cortex synaptic plasticity and behavior models of fear extinction. Considering the associative learning processes involved in contextual fear conditioning and extinction and the effect of DOI on the faster development of the later, it is plausible that activation of 5-HT_2A_R by this psychedelic accelerates context fear extinction likely in part through alterations in chromatin state at enhancer regions of genes predominantly involved in synapse organization and assembly such as *Mef2B*, *NeuroG2* and *Atoh1* – a gene network that may be located downstream of the genes showing transient transcription upon acute psychedelic administration. Recent observations have also reported that a putative non-hallucinogenic psychedelic analogue referred to as tabernanthalog promotes neural plasticity and reduces rodent behavior models relevant to depression (*65*). Further work will be needed to unravel the cellular signaling and neural target mechanism that may link psychedelic-induced synaptic plasticity and behavior, as well as the role, if any, of 5-HT_2A_R in these plasticity-related events induced by non-hallucinogenic 5-HT_2A_R agonist variants.

In conclusion, our study highlights the fundamental role of 5-HT_2A_R in the action of psychedelics and unveils persisting chromatin remodeling events following DOI administration linked to lasting synaptic plasticity and behavioral events. If generalizable to other psychedelics currently in clinical studies, these findings could also facilitate the understanding of psychopharmacological interventions whose mechanisms of action are not fully understood. Lastly, the overlap of the epigenomic markers of the action of the psychedelic DOI with loci associated with schizophrenia advocate for caution in the use of psychedelics in individuals at risk for psychosis.

## MATERIALS AND METHODS

### Drugs

DOI or 1-(2,5-dimethoxy-4-iodophenyl)-2-aminopropane was obtained from Sigma-Aldrich. Vehicle refers to saline (0.9%). All other chemicals were obtained from standard vendors.

### Animals

Experiments were performed on adult (10 – 20 weeks old) male mice randomly allocated into the different pre-treatment groups. Animals were housed on a 12 h light/dark cycle at 23 °C with food and water ad libitum, except during behavioral testing (always during the light cycle). All procedures were conducted in accordance with NIH guidelines and were approved by the Virginia Commonwealth University Animal Care and Use Committee. All efforts were made to minimize animal suffering and the number of animals used. Experiments involving dendritic spine structure, synaptic plasticity, and ChIP-seq/RNA-seq techniques were conducted in 129S6/SvEv mice (Taconic). Experiments involving behavior assays that required only the inclusion of wild-type animals were conducted in C57BL/6 mice (JAX). For fear acquisition and extinction assays, experiments were conducted in *5-HT_2A_R^+/+^* and *5-HT_2A_R^−/−^* mice that were offspring of heterozygote breeding(*15*). Breeders were backcrossed from 129S6/SvEv onto C57BL/6 for several generations (F5 was used for behavioral paradigms).

### Locomotor activity

Locomotor activity was monitored as previously described (*66*), with minor changes. Briefly, animals were injected with DOI (2 mg/kg, i.p.) or vehicle and immediately returned to their home cage. Evaluation of locomotor activity was conducted 24h after drug administration on a computerized three-dimensional activity monitoring system (Fusion v5.3; Omnitech Electronics Inc.) that interpolates the animal’s activity from interruption of infrared beams traversing the different planes of space. Mice were allowed to acclimate to the behavioral room for 1h prior to monitoring their individual locomotor activity on a commercial Plexiglas open field (25 × 25 × 20 cm) for 90 min. Horizontal activity is measured as the collapsed amount of beam interruptions in the *x*-*y* planes in 5 min periods. The open field was cleaned with 1% Roccal-D in between sessions.

### Light-dark boxes preference

Animals were injected with DOI (2 mg/kg, i.p.) or vehicle and immediately returned to their home cage. Preference for light or dark environment as a proxy for anxiety-like behaviors was assessed 18h after drug administration using a commercial open field in which space is equally divided into light and dark areas (25 × 12 × 20 cm each) connected through a small (5 × 5 cm) opening (*67*). The tracking system (Fusion v5.3; Omnitech Electronics Inc.) infers animal activity and position from interruptions to infrared beams traversing the different planes of space. After 1h acclimation to the behavioral room, mice were individually placed in the entrance to the dark compartment. The study monitored their preference for each area (light/dark) expressed as the percentage of time spent on each. The chambers were cleaned with 1% Roccal-D in between sessions. After completion of the experiment mice were returned to their home-cages.

### Forced-swimming test

The procedure was conducted 24h after DOI (2 mg/kg, i.p.) or vehicle administration (including a 1h acclimation to the behavioral room). 2 L (12.5 cm diameter × 21 cm height) beakers were 2/3-filled with water (~23°C). Mice were carefully introduced in the containers while being held by the tail so that the head would remain over the water line during the animal immersion. Sessions were recorded on digital video camera and immobility was scored during the last 4 min of the 6 min session by a trained analyst blind to the pre-treatment. Immobility was defined as passive floating with no additional activity except that needed to maintain the head of the animal above the water level (*68*).

### Novel object recognition

Assays were conducted as previously reported (*26, 69*), with minor modifications. Briefly, the testing arena consisted of an opaque rectangular plastic container with open top (32 × 20 × 23 cm). The objects employed (tissue culture flasks filled with wood chips and opaque white light bulbs) were of comparable volume and height (~10 cm) and were attached to the bottom of the container with double-side tape 10 cm away from the walls. Both the objects and chambers were cleaned thoroughly with diluted ethanol between tests to remove olfactory cues. Preference for a novel object — as opposed to a familiar one — was tested on mice that received DOI (2 mg/kg, i.p.) or vehicle 24h prior. Mice were allowed to get acclimate to the behavioral room for the duration of the whole session (at least 1h prior to the test). The test consisted of three stages, a 20 min habituation to the field, a 5 min acquisition trial in which two identical objects were present in the field and a 5 min recognition trial in which one of the identical objects had been replaced by a novel one. All stages were separated by 5 min during which the animals were returned to their home cage. Object exploration was defined as the animal licking, sniffing or touching the object with the forepaws while sniffing. Climbing an object was not considered in the exploration time. The exploration time for each side (same object on both sides) during the acquisition and the exploration time for the novel object during the recognition was assessed on digital video camera recordings by a trained analyst blind to the pre-treatment. The exploratory preference for the novel object was calculated as percentage of total exploratory time spent exploring the novel object during the recognition stage. Place preference was controlled during the acquisition stage and by alternating the position of the novel object between left and right during the recognition stage.

### Prepulse inhibition of startle

Assays were conducted as previously reported (*70*) with minor modifications. Briefly, recording of the startle magnitude in response to acoustic stimuli was performed using the SR-LAB Startle Response System (San Diego Instruments). Mice that had received DOI (2 mg/kg) or vehicle 24h prior were presented with a startling stimulus of 119 dB (20 ms) preceded (80 ms interstimulus interval), or not, by (20 ms) prepulses of 73 dB, 77 dB and 85 dB (20 ms). Background noise was set to 69 dB. Mice were habituated to the testing room at least 1h prior to the start of the experiment. The experiment proceeded as follows: Animals were placed in the mouse-sized startle chambers and allowed to acclimate to chamber and background noise for 5 min. Mice were presented with 5 stimulus-only trials at the beginning and the end of the session (used for control, but not included in the analysis). During the session, animals were randomly subjected to 65 trials in five different categories (13 of each): stimulus-only, no stimulus, 73 dB prepulse and stimulus, 77 dB prepulse and stimulus, 85 dB prepulse and stimulus. Each session lasted about a total of 30 min. Prepulse inhibition of startle (PPI) for each prepulse and trial was determined as percentage of the startle amplitude (V_max_) of the prepulse+stimulus trial relative to the average of the startle amplitude (V_max_) for the 13 stimulus-only trials.

### Contextual fear acquisition and extinction

Fear conditioning tests were conducted using two commercially supplied Near Infrared Video Freeze Systems controlling each testing chamber (MED-VFCNIR-M, Med Associates, Inc., St. Albans, VT), each operating four test chambers. Each polycarbonate test chamber (32 × 25 × 25 cm) was equipped with a metal grid floor and was enclosed in a sound-attenuated box (63.5 × 35.5 × 76 cm). Video images at 30 fps (640 × 480 pixels, down-sampled to ~1 pixel per visible mm^2^) were collected. A freezing event was automatically designated if the absence of movement was 1 s or longer (*i.e*., negligible variation in the pixel composition in between frames for 30 consecutive frames). Mice were acclimated to the behavioral testing room (with white noise in the background) for at least 1h in the 2 days prior to experimental sessions. White noise was present at all times when the animals were held in the testing room. Mice were handled for 2 min (each) on the 2 days prior to the start of the testing. Two different contexts (A and B) with different spatial, visual and olfactory cues were employed. Context A was composed of a standard polycarbonate squared cage (see dimensions above) with metallic walls and shock-delivering grid on the floor, with visible and IR illumination, and with lemon scent (artificial lemon flood flavoring) on the bedding. Context B was a modified chamber with a black triangular adapter, white opaque plastic continuous floor, without visible light but with IR light and cardamom scent (cardamom artificial food flavoring) on the bedding. Experimental sessions were performed on two consecutive days. On day 1, fear-acquisition was established in Context A. The acquisition stage period conditioned the mice to associate the context (A) with a noxious stimulus (foot-shocks). After a 90s stimulus-free period at the beginning of the experiment (baseline), mice received 6 unconditioned stimuli consisting of 2s scrambled 0.70 mA foot-shocks delivered through the grid floor separated by 90 s inter-stimulus intervals. Freezing time was measured as described above during the baseline and during the inter-stimulus intervals. For all other periods, freezing time was measured in 5 min units. Animals were returned to their home cages immediately after completion of the 90 s period following the last foot-shock. After a 2h holding period, the expression stage followed. Animals were placed in Context A for 5 min. During this time, foot-shocks were not delivered. This period served as a control for conditioning and preliminary extinction training (noxious stimulus-free Context A). Depending on testing conditions, DOI (2 mg/kg, i.p.) or vehicle was administered 10 min after the expression stage or 24h prior to the acquisition stage. At the end of the experimental session animals were returned to the vivarium. On day 2, the generalization of the fear response was evaluated in Context B for 5 min. The purpose of this test was to evaluate and quantify the expression of generalization of fear in a novel context (*i.e*., expression of freezing behavior in a novel environment). Mice were held for 2 additional hours in their home cage prior to the extinction test. During extinction training, mice were returned to Context A for 20 min and observed for freezing behavior over time in periods of 5 min in the absence of any aversive stimuli.

### Dendritic spine analysis

Dendritic spine analysis assays were carried out as previously reported (*26*), with minor modifications. Briefly, adeno-associated virus (AAV) serotype 8 expressing eYFP under the *CaMKIIα* promoter was produced at the University of North Carolina at Chapel Hill Vector Core. Mice were anesthetized with isoflurane during the surgery and perfusion. The virus was delivered bilaterally with a Hamilton syringe at a rate of 0.1 μl/min for a total volume of 0.5 μl on each side. The following coordinates were used: +1.6 mm rostrocaudal, −2.4 mm dorsoventral, and +2.6 mm mediolateral from bregma (relative to dura) with a 10°C lateral angle. Previous findings have validated expression of eYFP in CaMKIIα-positive neurons, but not in parvalbumin-positive neurons in the frontal cortex (*26*). All experiments were performed at least three weeks after surgery, when transgene expression is maximal (*26*). For spine analysis, apical dendritic segments 50-150 μm away from the soma were randomly chosen from AAV-infected cells that express eYFP. Images were acquired from a 4% paraformaldehyde-fixed 50 μm coronal slice, using a confocal fluorescence microscope (CLSM710, Carl Zeiss). CaMKIIα-positive pyramidal neurons in L5 expressing eYFP were confirmed by their characteristic triangular somal shape. To qualify for spine analysis, dendritic segments had to satisfy the following requirements: (i) the segment had to be completely filled (all endings were excluded), (ii) the segment must have been at least 50 μm from the soma, and (iii) the segment could not be overlapping with other dendritic branches. Dendritic segments were imaged using a ×63 lens (numerical aperture 1.46; Carl Zeiss) and a zoom of 2.5. Pixel size was 0.09 μm in the x-y plane and 0.2 μm in the z plane. Images were taken with a resolution of 1024 × 300 (the y dimension was adjusted to the particular dendritic segment to expedite imaging), the pixel dwell time was 1.27 μm/s and the line average was set to 4. Only one dendrite per neuron on 4-5 neurons per animal per experimental group was analyzed. For quantitative analysis of spine size and shape, NeuronStudio was used with the rayburst algorithm described previously (*26*). NeuronStudio classifies spines as stubby, thin or mushroom on the basis of the following values: (i) aspect ratio, (ii) head-to-neck ratio and (iii) head diameter. Spines with a neck can be classified as either thin or mushroom, and those without a neck are classified as stubby. Spines with a neck are labelled as thin or mushroom on the basis of head diameter. These parameters have been verified by comparison with trained human operators fully blinded across groups.

### Whole-cell patch-clamp recordings and LTP induction

Whole-cell patch clamp recordings from neurons in layers 2/3 (L2/3) of somatosensory cortex were obtained from 8-10-week-old male mice as previously described (*26*). Prior to recording, mice were housed in pairs, with one mouse of each pair designated an “experimental” mouse and the other mouse designated a “companion”. Twenty-four hours prior to recording, experimental mice of each pair received a single injection (i.p.) of either saline (controls, n=5) or DOI (2,5-Dimethoxy-4-iodoamphetamine hydrochloride; 2 mg/kg, n= 5). The investigator performing the recordings was blind to treatment condition. Companion mice, who received a single saline injection at the same time as experimental mice, were used to match housing/social conditions across control and DOI experimental animals in the 24 hr period prior to recording, but were then discarded. Experimental mice were deeply anesthetized with isoflurane and decapitated. Brains were rapidly removed and submerged in chilled (~4°C) sucrose-artificial cerebrospinal fluid (sucrose-aCSF; 233.7 mM sucrose, 26 mM NaHCO_3_, 3 mM KCl, 8 mM MgCl_2_, 0.5 mM CaCl_2_, 20 mM glucose and 0.4 mM ascorbic acid) that was continuously bubbled with carbogen (95% O_2_–5% CO_2_). Acute coronal slices (350 μm-thick) were sectioned on a VT1000S vibratome (Leica) and transferred to a recovery chamber bubbled with carbogen and aCSF (117 mM NaCl, 4.7 mM KCl, 1.2 mM MgSO_4_, 2.5 mM CaCl_2_, 1.2 mM NaH_2_PO_4_, 24.9 mM NaHCO_3_ and 11.5 mM glucose) for 1h at room temperature. Slices were transferred to the recording chamber and perfused with 31°C oxygenated aCSF containing the GABA_A_ receptor antagonist gabazine (10 μM). Whole-cell electrodes were pulled from borosilicate glass (pipette resistance 3–4 MΩ) and filled with (in mM): 120 Cs-methanesulfonate, 10 HEPES, 0.5 EGTA, 8 NaCl, 5 TEA-CL, 4 Mg-ATP, 0.4 NaGTP, and 10 phosphocreatine. Internal solution was adjusted to 280–290 mOsm and 7.3 pH. L2/3 neurons chosen for recording had pyramidal-shaped somata. Following formation of a gigaseal, the neuronal membrane was ruptured and set to a holding potential of - 70 mV and allowed to equilibrate for ~3 mins. Subsequent recordings were carried out in voltage-clamp mode with a Multiclamp 700B amplifier (Molecular Devices). Analog signals were low-pass filtered at 2 kHz, digitized at 5 kHz and analyzed with pClamp 10 software (Molecular Devices). LTP of L4 to L2/3 synapses was elicited using a pairing-protocol in which presynaptic stimulation of L4 was paired with brief postsynaptic depolarization of L2/3 neurons situated within the same column as described previously (*26*). A tungsten concentric bipolar electrode was placed in L4 and used to evoke excitatory postsynaptic currents (EPSCs) in overlying L2/3 neurons at 0.1 Hz through the entire experiment. EPSCs evoked by L4 stimulation were recorded at −70 mV for a baseline period of ~3-5 mins, followed by brief membrane depolarization to 0 mV for 10 mins, then the holding potential was returned to −70mV for the duration of 35-40 mins. Series resistance was measured at the beginning and end of each recording. Any neuron showing >30% change was discarded from the experiment. Offline analysis was conducted with Clampfit software (Molecular Devices). Mean EPSC amplitudes were normalized to their respective averaged baseline values. Data are plotted as percent change from averaged baseline values.

### Nuclei isolation and sorting via FACS

Nuclei isolation was conducted using a published protocol (*30, 71*). All steps were conducted on ice, and all centrifugation was conducted at 4 °C. One piece of mouse frontal cortex tissue (6-10 mg) was placed in 3 ml of ice-cold nuclei extraction buffer (NEB) [0.32 M sucrose, 5 mM CaCl_2_, 3 mM Mg(Ac)_2_, 0.1 mM EDTA, 10 mM tris-HCl, and 0.1%(v/v) Trition X-100] with 30 μl of freshly added protease inhibitor cocktail (PIC, Sigma-Aldrich), 3 μl of 100 mM phenylmethylsulfonyl fluoride (PMSF, Sigma-Aldrich) in Isopropyl alcohol, 3 μl of 1 M dithiothreitol (DTT, Sigma-Aldrich), and 4.5 μl of ribonuclease (RNase) inhibitor (2313A,Takara Bio). The tissue was homogenized in a tissue grinder (D9063, Sigma-Aldrich). The homogenate was filtered with a 40 μm cell strainer (22-363-547, Thermo Fisher Scientific) and collected in a 15-ml centrifuge tube. The cell suspension was centrifuged at 1000 RCF for 10 min. The supernatant was discarded, and the pellet was resuspended in 0.5 ml of ice-cold NEB with 5 μl of freshly added PIC, 0.5 μl of PMSF, 0.5 μl of DTT, and 0.75 μl of RNase inhibitor. 500 μl of the sample was mixed with 750 μl of 50%(w/v) iodixanol (made by mixing 4 ml of OptiPrep™ gradient (Sigma-Aldrich) and 0.8 ml of diluent [150 mM KCl, 30 mM MgCl_2_, and 120 mM tris-HCl]). The mixture was centrifuged at 10,000 RCF for 20 min. Then, the supernatant was removed and 300 μl of 2%(w/v) normal goat serum (50062Z, Life technologies) in Dulbecco’s PBS (DPBS, Life technologies) was added to resuspend the nuclei pellet. To separate NeuN^+^ and NeuN^−^ fractions, 6 μl of 2 ng/μl anti-NeuN antibody conjugated with Alexa 488 (MAB377X, Millipore) in DPBS was added into the nuclei suspension. The suspension was mixed well and incubated at 4 °C for 1 h on a rotator mixer (Labnet). After incubation, the sample was sorted into NeuN^+^ and NeuN^−^ fractions using a BD FACSAria™ cell sorter (BD Biosciences). The sorted NeuN^+^ nuclei were directly used for RNA-seq experiment. 200 μl of sorted NeuN^+^ nuclei suspension, containing ~26,000 nuclei, was added into 800 μl of ice-cold PBS for ChIP-seq experiment. 200 μl of 1.8 M sucrose solution, 10 μl of 1 M CaCl_2_, and 3 μl of 1 M Mg(Ac)_2_ were added into the mixture. The solution was mixed well and incubated on ice for 15 min. Then, the sample was centrifuged at 1800 RCF at 4 °C for 15 min. The supernatant was discarded and the pellet was resuspended in 60 μl of PBS with 0.6 μl of freshly added PIC and 0.6 μl of PMSF and stored on ice until use for ChIP-seq.

### Construction of ChIP-seq libraries

~10,000 NeuN^+^ nuclei were used to produce each ChIP-seq library and two technical replicates were generated for each brain sample. One input per sample was generated using ~4,000 nuclei. ChIP was carried out using multiplexed MOWChIP-seq with MNase digestion for chromatin fragmentation and anti-H3K27ac (39135, Active Motif) antibody, following a published protocol (*30*).

### Construction of RNA-seq libraries

~5000 NeuN^+^ nuclei were used to produce each RNA-seq library and two technical replicates were generated for each brain sample. RNA extraction from 50 μl of nuclei suspension from FACS (containing 5000 sorted NeuN^+^ nuclei) was conducted using RNeasy Mini Kit (74104, Qiagen) and RNase-Free DNase Set (79254, Qiagen), following the manufacturer’s instruction. Extracted mRNA in 30-μl volume was concentrated by ethanol precipitation and resuspended in 4.6 μl of RNase-free water. cDNA was prepared using the SMART-seq2 protocol (*31*) with minor modification. ~2 ng of mRNA in 4.6 μl of water was mixed with 2 μl of 100 μM oligo-dT primer and 2 μl of 10 mM dNTP mix. After being denatured at 72 °C for 3 min, the mRNA solution was immediately placed on ice. 11.4 μl of reverse transcript mix [made from 1 μl of SuperScript II reverse transcriptase (200 U/μl), 0.5 μl of RNAse inhibitor (40 U/μl), 4 μl of Superscript II first-strand buffer, 1 μl of DTT (100mM), 4 μl of 5 M Betaine, 0.12 μl of 1 M MgCl_2_, 0.2 μl of TSO (100 μM), 0.58 μl of nuclease-free water] was mixed with the mRNA solution and the mixture was incubated at 42 °C for 90 min, followed by 10 cycles of (50 °C for 2 min, 42 °C for 2 min). The reaction was finally inactivated at 70 °C for 15 min. 20 μl of first-strand mixture was then mixed with 25 μl of KAPA HiFi HotStart ReadyMix, 0.5 μl of (100 μM) IS PCR primers, 0.5 μl of Evagreen dye, and 4 μl of nuclease-free water. Generated cDNA was amplified by incubation at 98 °C for 1 min, followed by 9-11 cycles of (98 °C 15 s, 67 °C 30 s, 72 °C 6 min). After PCR amplification, 50 μl of PCR mix was purified by using 50 μl of SPRIselect beads. ~600 pg of purified cDNA was used to produce a RNA-seq library using Nextera XT DNA Library Preparation kit (FC-131-1024, Illumina), following the manufacturer instructions.

### Sequencing

The fragment size of ChIP-seq and RNA-seq libraries was measured using high sensitivity DNA analysis kit (5067-4626, Agilent) on a TapeStation system (2200, Agilent). The concentration of each library was examined using a KAPA library quantification kit (KK4809, Kapa Biosystems), and then the quantified libraries were pooled at 10 nM. The libraries were sequenced by Illumina HiSeq 4000 with single-end 50-nt read. Around 15 million reads were generated for each ChIP-seq library, 10 million reads for each input library, and 11 million reads for each RNA-seq library.

### ChIP-seq reads alignment and normalization

Sequencing reads were trimmed using default settings by Trim Galore! (Babraham Institute). Trimmed reads were aligned to the mm10 genome with Bowtie2 (*72*). Peaks were called using MACS2 (*73*) (q < 0.05). Blacklisted regions in the mm10 as defined by ENCODE were removed to improve data quality. Mapped reads from ChIP and input samples were extended by 100 bp on either side (250 bp total) and a normalized signal was calculated.

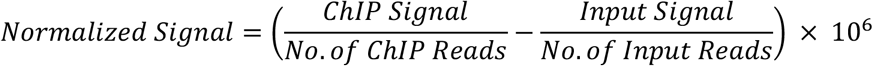

For visualization in IGV (Broad Institute), the signal was calculated in 100bp windows over the entire genome and output as a bigWig file.

### Differential enhancers

We first predicted enhancer regions (1000 bp in width with peak summit +/-500 bp) by identifying H3K27ac peaks that did not intersect with promoters (defined as TSS +/- 2000 bp) (*29*). Consensus enhancer sets were generated for each experimental group (VEH, 24 h, 48 h and 7d) using Diffbind (*74*) and then combined to generate an overall consensus enhancer set. In the process, a majority rule was applied to identify the consensus enhancers when all technical/biological replicates were considered. For example, among all 12 replicates of H3K27ac data sets in one experimental group, a consensus enhancer must be present in at least 7. Differential enhancers between any two experimental groups were identified from the overall consensus enhancer set using the bioconductor DESeq2 package (*75*) (FDR<0.05) with Benjamini-Hochberg method (*76*).

### K-means clustering, GO term analysis, TF motif analysis

We also performed K-means clustering on differential enhancers and DEGs across experimental groups. In each row, the values (x) were normalized by dividing by 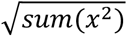. The optimal number of clusters was determined using Silhouette method (*77*). We created a list of the enhancer target genes by examining the correlation between gene expression (as reflected by RNA-seq data) and H3K27ac intensity at the candidate enhancers (*78*). Compared to assigning an enhancer to its nearest gene, this approach produces a much higher percentage of enhancer-gene pairs that match the results determined experimentally using techniques including Hi-C (*78*). All the enhancers that were not linked with their target genes after the step were then processed using ChIPSeeker (*79*) with default condition and TxDb.Mmusculus.UCSC.mm10.knownGene to link to their nearest genes (as target genes). All the target genes identified, either by correlation between H3K27ac and RNA-seq or by linking with nearest genes, were combined for downstream analysis. GO biological processes enrichment test was performed using the bioconductor clusterProfiler (*80*) package (with FDR adjusted p-value < 0.01). Motif analysis was also performed to determine enriched transcription factor binding motifs among the enhancer regions using Homer (*81*) (with options –size 200 –p 16, and q-value < 0.01).

### GWAS loci association

Differential peak locations were converted from mm10 to hg19 through UCSC’s liftOver tool. GWAS datasets were obtained from NHGRI-EBI’s GWAS Catalog (https://www.ebi.ac.uk/gwas/home). SNPs that did not have a reference SNP ID were removed. Only SNPs present in the 1000 Genome Project Phase 3 European population were kept. For each GWAS set, if there were *n* SNPs in the GWAS set, then *n* SNPs would be randomly selected from all SNPs present in the 1000 Genome European population set, retaining an identical distribution of SNPs across the chromosomes. The number of differential peaks that overlapped with this SNP test set was calculated and repeated for a total of 10,000 runs (Monte Carlo randomization). From this distribution, we used the R function pnorm, in conjunction with the actual overlap of the GWAS set with the differential peaks, to calculate a p-value.

### RNA-seq data analysis

Sequencing reads were trimmed under default settings using Trim Galore!. The trimmed reads were mapped to GRCm38 genome by hisat2 (*82*). Mapped bam files were imported into SeqMonk v1.47.1 (Babraham Institute) and data sets with poor quality (percentage of reads aligned to exons <40%) were discarded. featureCounts (*83*) was used to count reads and DEGs were determined by pairwise comparison using DESeq2 (FDR<0.05).

### Weighted gene co-expression network analysis

A signed co-expression network was built using the WGCNA package in R (*36*). We firstly normalized gene expression values using log2(FPKM) (Fragments Per Kilobase per Million mapped reads) and then genes were ranked based on their median absolute deviation (MAD). The genes with over 0.3 at MAD were selected as input for the network construction. We converted the normalized gene expression values into an adjacency matrix using the function *a*_*ij*_ = |*cor*(*x*_*i*_, *x*_*j*_)|^*β*^, where *x*_*i*_and *x*_*j*_ were the expression data of two genes. The value of β was determined to be 7 using the approximate scale-free topology (*84*). We further performed the topological overlap measure to transform the adjacency matrix into a topological overlap matrix using the blockwiseModules function in the package (with parameters corType = “bicor”, TOMType = “signed”, minModuleSize = 30, reassignThreshold = 0, mergeCutHeight = 0.15). Modules were then detected based on the topological overlap matrix and the gene expression level of each module was characterized by the module eigengene. The top 100 intramodular hub genes (the most highly connected genes within the module) were identified using the softConnedticity function in the package. The relationship between these hub genes were visualized using Cytoscape (*85*).

### Statistical analysis

All statistical analysis was performed with GraphPad Prism software version 9. Microscope images were taken by an observer who was blind to the experimental conditions. No statistical methods were used to predetermine sample sizes, but our sample sizes are similar to those reported in our previous publications (*26*). Data distribution was assumed to be normal, but this was not formally tested. Animals were randomly allocated into the different experimental groups. Datapoints were excluded based on previously established criterion and were set to ± 2 s.d. from the group mean. Statistical significance of experiments involving three or more groups and two or more treatments was assessed by two-way ANOVA followed by Sidak’s multiple comparison test. Statistical significance of experiments involving time courses and two treatments was assessed by two-way repeated measures ANOVA. Statistical significance of experiments involving three or more groups was assessed by one-way ANOVA followed by Sidak’s multiple comparison test. Statistical significance of experiments involving two groups was assessed by Student’s *t* test. The level of significance was set at p = 0.05. All values in the figure legends represent mean ± s.e.m.

## Acknowledgments

We thank Jennifer Jimenez and Yashu Sampathkumar for technical assistance with behavior assays.

## Author contributions

M.d.l.F.R., B.Z., C.L. and J.G.-M. designed experiments, analyzed data and wrote the manuscript. C.L. and J.G.-M. supervised the research and obtained funding. M.d.l.F.R., J.M.S., R.T. and S.S., supervised by J.G.-M., performed behavioral and synaptic structure assays. B.Z., Z.Z. and L.B.N., supervised by C.L., performed ChIP-seq and RNA-seq assays and conducted bioinformatic analysis. C.A.G., supervised by G.W.H., performed electrophysiological studies. J.T.W. and P.M.B. provided advice on behavioral assays and editorial suggestions on early drafts of the report. All authors discussed the results and commented on the manuscript prior to submission for publication consideration.

## Data and materials availability

All data needed to evaluate the conclusions in the paper are present in the paper and/or Supplementary Materials. ChIP-seq and RNA-seq data sets are deposited in the Gene Expression Omnibus (GEO) repository with the following accession number: GSE161626. The following secure token has been created to allow review of record GSE161626 while it remains in private status: utobaegezfurbol

## References

1. V. Krishnan, E. J. Nestler, The molecular neurobiology of depression. Nature 455, 894–902 (2008).

2. R. C. Kessler, P. Berglund, O. Demler, R. Jin, D. Koretz, K. R. Merikangas, A. J. Rush, E. E. Walters, P. S. Wang, R. National Comorbidity Survey, The epidemiology of major depressive disorder: results from the National Comorbidity Survey Replication (NCS-R). JAMA 289, 3095–3105 (2003).

3. R. S. Duman, G. K. Aghajanian, Synaptic dysfunction in depression: potential therapeutic targets. Science 338, 68–72 (2012).

4. T. R. Insel, P. S. Wang, The STAR*D trial: revealing the need for better treatments. Psychiatr Serv 60, 1466–1467 (2009).

5. R. A. Glennon, Classical hallucinogens: an introductory overview. NIDA Res Monogr 146, 4–32 (1994).

6. D. E. Nichols, Psychedelics. Pharmacol Rev 68, 264–355 (2016).

7. J. B. Hanks, J. Gonzalez-Maeso, Animal models of serotonergic psychedelics. ACS Chem Neurosci 4, 33–42 (2013).

8. R. L. Carhart-Harris, M. Bolstridge, J. Rucker, C. M. Day, D. Erritzoe, M. Kaelen, M. Bloomfield, J. A. Rickard, B. Forbes, A. Feilding, D. Taylor, S. Pilling, V. H. Curran, D. J. Nutt, Psilocybin with psychological support for treatment-resistant depression: an open-label feasibility study. Lancet Psychiatry 3, 619–627 (2016).

9. R. L. Carhart-Harris, M. Bolstridge, C. M. J. Day, J. Rucker, R. Watts, D. E. Erritzoe, M. Kaelen, B. Giribaldi, M. Bloomfield, S. Pilling, J. A. Rickard, B. Forbes, A. Feilding, D. Taylor, H. V. Curran, D. J. Nutt, Psilocybin with psychological support for treatment-resistant depression: six-month follow-up. Psychopharmacology (Berl) 235, 399–408 (2018).

10. A. K. Davis, F. S. Barrett, D. G. May, M. P. Cosimano, N. D. Sepeda, M. W. Johnson, P. H. Finan, R. R. Griffiths, Effects of Psilocybin-Assisted Therapy on Major Depressive Disorder: A Randomized Clinical Trial. JAMA Psychiatry (2020).

11. R. R. Griffiths, M. W. Johnson, M. A. Carducci, A. Umbricht, W. A. Richards, B. D. Richards, M. P. Cosimano, M. A. Klinedinst, Psilocybin produces substantial and sustained decreases in depression and anxiety in patients with life-threatening cancer: A randomized double-blind trial. J Psychopharmacol 30, 1181–1197 (2016).

12. M. Hibicke, A. N. Landry, H. M. Kramer, Z. K. Talman, C. D. Nichols, Psychedelics, but Not Ketamine, Produce Persistent Antidepressant-like Effects in a Rodent Experimental System for the Study of Depression. ACS Chem Neurosci 11, 864–871 (2020).

13. L. P. Cameron, C. J. Benson, L. E. Dunlap, D. E. Olson, Effects of N, N-Dimethyltryptamine on Rat Behaviors Relevant to Anxiety and Depression. ACS Chem Neurosci (2018).

14. T. S. Ray, Psychedelics and the human receptorome. PloS one 5, e9019–e9019 (2010).

15. J. Gonzalez-Maeso, N. V. Weisstaub, M. Zhou, P. Chan, L. Ivic, R. Ang, A. Lira, M. Bradley-Moore, Y. Ge, Q. Zhou, S. C. Sealfon, J. A. Gingrich, Hallucinogens Recruit Specific Cortical 5-HT(2A) Receptor-Mediated Signaling Pathways to Affect Behavior. Neuron 53, 439–452 (2007).

16. C. E. Canal, Head-twitch response in rodents induced by the hallucinogen 2, 5-dimethoxy-4-iodoamphetamine: a comprehensive history, a re-evaluation of mechanisms, and its …. Drug Testing and Analysis (2012).

17. M. de la Fuente Revenga, J. M. Shin, H. Z. Vohra, K. S. Hideshima, M. Schneck, J. L. Poklis, J. Gonzalez-Maeso, Fully automated head-twitch detection system for the study of 5-HT2A receptor pharmacology in vivo. Sci Rep 9, 14247 (2019).

18. P. Tovote, J. P. Fadok, A. Luthi, Neuronal circuits for fear and anxiety. Nat Rev Neurosci 16, 317–331 (2015).

19. N. Spruston, Pyramidal neurons: dendritic structure and synaptic integration. Nat Rev Neurosci 9, 206–221 (2008).

20. A. J. Koleske, Molecular mechanisms of dendrite stability. Nat Rev Neurosci 14, 536–550 (2013).

21. N. K. Savalia, L. X. Shao, A. C. Kwan, A Dendrite-Focused Framework for Understanding the Actions of Ketamine and Psychedelics. Trends Neurosci (2020).

22. C. Ly, A. C. Greb, L. P. Cameron, J. M. Wong, E. V. Barragan, P. C. Wilson, K. F. Burbach, S. Soltanzadeh Zarandi, A. Sood, M. R. Paddy, W. C. Duim, M. Y. Dennis, A. K. McAllister, K. M. Ori-McKenney, J. A. Gray, D. E. Olson, Psychedelics Promote Structural and Functional Neural Plasticity. Cell Rep 23, 3170–3182 (2018).

23. R. T. Roppongi, N. Kojima, K. Hanamura, H. Yamazaki, T. Shirao, Selective reduction of drebrin and actin in dendritic spines of hippocampal neurons by activation of 5-HT(2A) receptors. Neurosci Lett 547, 76–81 (2013).

24. M. Kondoh, T. Shiga, N. Okado, Regulation of dendrite formation of Purkinje cells by serotonin through serotonin1A and serotonin2A receptors in culture. Neurosci Res 48, 101–109 (2004).

25. D. J. Christoffel, S. A. Golden, M. Heshmati, A. Graham, S. Birnbaum, R. L. Neve, G. E. Hodes, S. J. Russo, Effects of Inhibitor of kappaB Kinase Activity in the Nucleus Accumbens on Emotional Behavior. Neuropsychopharmacology (2012).

26. D. Ibi, M. de la Fuente Revenga, N. Kezunovic, C. Muguruza, J. M. Saunders, S. A. Gaitonde, J. L. Moreno, M. K. Ijaz, V. Santosh, A. Kozlenkov, T. Holloway, J. Seto, A. Garcia-Bea, M. Kurita, G. E. Mosley, Y. Jiang, D. J. Christoffel, L. F. Callado, S. J. Russo, S. Dracheva, J. F. Lopez-Gimenez, Y. Ge, C. R. Escalante, J. J. Meana, S. Akbarian, G. W. Huntley, J. Gonzalez-Maeso, Antipsychotic-induced Hdac2 transcription via NF-kappaB leads to synaptic and cognitive side effects. Nat Neurosci 20, 1247–1259 (2017).

27. J. Gonzalez-Maeso, T. Yuen, B. J. Ebersole, E. Wurmbach, A. Lira, M. Zhou, N. Weisstaub, R. Hen, J. A. Gingrich, S. C. Sealfon, Transcriptome fingerprints distinguish hallucinogenic and nonhallucinogenic 5-hydroxytryptamine 2A receptor agonist effects in mouse somatosensory cortex. J Neurosci 23, 8836–8843 (2003).

28. C. D. Nichols, E. Sanders-Bush, A single dose of lysergic acid diethylamide influences gene expression patterns within the mammalian brain. Neuropsychopharmacology 26, 634–642 (2002).

29. Z. Cao, C. Chen, B. He, K. Tan, C. Lu, A microfluidic device for epigenomic profiling using 100 cells. Nat Methods 12, 959–962 (2015).

30. B. Zhu, Y. P. Hsieh, T. W. Murphy, Q. Zhang, L. B. Naler, C. Lu, MOWChIP-seq for low-input and multiplexed profiling of genome-wide histone modifications. Nat Protoc 14, 3366–3394 (2019).

31. S. Picelli, A. K. Bjorklund, O. R. Faridani, S. Sagasser, G. Winberg, R. Sandberg, Smart-seq2 for sensitive full-length transcriptome profiling in single cells. Nat Methods 10, 1096–1098 (2013).

32. S. Picelli, O. R. Faridani, A. K. Bjorklund, G. Winberg, S. Sagasser, R. Sandberg, Full-length RNA-seq from single cells using Smart-seq2. Nat Protoc 9, 171–181 (2014).

33. A. S. Nord, A. E. West, Neurobiological functions of transcriptional enhancers. Nat Neurosci 23, 5–14 (2020).

34. S. Venteo, S. Desiderio, P. Cabochette, A. Deslys, P. Carroll, A. Pattyn, Neurog2 Deficiency Uncovers a Critical Period of Cell Fate Plasticity and Vulnerability among Neural-Crest-Derived Somatosensory Progenitors. Cell Rep 29, 2953–2960 e2952 (2019).

35. S. Mitsui, S. Yamaguchi, T. Matsuo, Y. Ishida, H. Okamura, Antagonistic role of E4BP4 and PAR proteins in the circadian oscillatory mechanism. Genes Dev 15, 995–1006 (2001).

36. P. Langfelder, S. Horvath, WGCNA: an R package for weighted correlation network analysis. BMC Bioinformatics 9, 559 (2008).

37. L. Wang, Y. Wang, C. Duan, Q. Yang, Inositol phosphatase INPP4A inhibits the apoptosis of in vitro neurons with characteristic of intractable epilepsy by reducing intracellular Ca(2+) concentration. Int J Clin Exp Pathol 11, 1999–2007 (2018).

38. P. G. Zagnoni, C. Albano, Psychostimulants and epilepsy. Epilepsia 43 Suppl 2, 28–31 (2002).

39. F. Nau, Jr., J. Miller, J. Saravia, T. Ahlert, B. Yu, K. I. Happel, S. A. Cormier, C. D. Nichols, Serotonin 5-HT(2) receptor activation prevents allergic asthma in a mouse model. Am J Physiol Lung Cell Mol Physiol 308, L191–198 (2015).

40. A. Klingseisen, A. M. Ristoiu, L. Kegel, D. L. Sherman, M. Rubio-Brotons, R. G. Almeida, S. Koudelka, S. K. Benito-Kwiecinski, R. J. Poole, P. J. Brophy, D. A. Lyons, Oligodendrocyte Neurofascin Independently Regulates Both Myelin Targeting and Sheath Growth in the CNS. Dev Cell 51, 730–744 e736 (2019).

41. J. C. McIntyre, W. B. Titlow, T. S. McClintock, Axon growth and guidance genes identify nascent, immature, and mature olfactory sensory neurons. J Neurosci Res 88, 3243–3256 (2010).

42. B. Zonta, S. Tait, S. Melrose, H. Anderson, S. Harroch, J. Higginson, D. L. Sherman, P. J. Brophy, Glial and neuronal isoforms of Neurofascin have distinct roles in the assembly of nodes of Ranvier in the central nervous system. J Cell Biol 181, 1169–1177 (2008).

43. O. Penagarikano, D. H. Geschwind, What does CNTNAP2 reveal about autism spectrum disorder? Trends Mol Med 18, 156–163 (2012).

44. P. Rodenas-Cuadrado, N. Pietrafusa, T. Francavilla, A. La Neve, P. Striano, S. C. Vernes, Characterisation of CASPR2 deficiency disorder--a syndrome involving autism, epilepsy and language impairment. BMC Med Genet 17, 8 (2016).

45. J. I. Friedman, T. Vrijenhoek, S. Markx, I. M. Janssen, W. A. van der Vliet, B. H. Faas, N. V. Knoers, W. Cahn, R. S. Kahn, L. Edelmann, K. L. Davis, J. M. Silverman, H. G. Brunner, A. G. van Kessel, C. Wijmenga, R. A. Ophoff, J. A. Veltman, CNTNAP2 gene dosage variation is associated with schizophrenia and epilepsy. Mol Psychiatry 13, 261–266 (2008).

46. A. J. Majmundar, W. J. Wong, M. C. Simon, Hypoxia-inducible factors and the response to hypoxic stress. Mol Cell 40, 294–309 (2010).

47. T. R. Anju, C. S. Paulose, Amelioration of hypoxia-induced striatal 5-HT(2A) receptor, 5-HT transporter and HIF1 alterations by glucose, oxygen and epinephrine in neonatal rats. Neurosci Lett 502, 129–132 (2011).

48. D. E. Olson, The Subjective Effects of Psychedelics May Not Be Necessary for Their Enduring Therapeutic Effects. ACS Pharmacol Transl Sci (2020).

49. D. B. Yaden, R. R. Griffiths, The Subjective Effects of Psychedelics Are Necessary for Their Enduring Therapeutic Effects. ACS Pharmacol Transl Sci (2020).

50. F. X. Vollenweider, M. F. Vollenweider-Scherpenhuyzen, A. Babler, H. Vogel, D. Hell, Psilocybin induces schizophrenia-like psychosis in humans via a serotonin-2 agonist action. Neuroreport 9, 3897–3902 (1998).

51. Y. Schmid, F. Enzler, P. Gasser, E. Grouzmann, K. H. Preller, F. X. Vollenweider, R. Brenneisen, F. Muller, S. Borgwardt, M. E. Liechti, Acute Effects of Lysergic Acid Diethylamide in Healthy Subjects. Biol Psychiatry (2015).

52. R. Griffiths, W. Richards, M. Johnson, U. McCann, R. Jesse, Mystical-type experiences occasioned by psilocybin mediate the attribution of personal meaning and spiritual significance 14 months later. J Psychopharmacol 22, 621–632 (2008).

53. L. S. Kaertner, M. B. Steinborn, H. Kettner, M. J. Spriggs, L. Roseman, T. Buchborn, M. Balaet, C. Timmermann, D. Erritzoe, R. L. Carhart-Harris, Positive expectations predict improved mental-health outcomes linked to psychedelic microdosing. Sci Rep 11, 1941 (2021).

54. R. M. Bastle, I. Maze, Chromatin Regulation in Complex Brain Disorders. Curr Opin Behav Sci 25, 57–65 (2019).

55. J. Graff, L. H. Tsai, Histone acetylation: molecular mnemonics on the chromatin. Nat Rev Neurosci 14, 97–111 (2013).

56. D. E. Babinski, K. A. Neely, D. M. Ba, G. Liu, Depression and Suicidal Behavior in Young Adult Men and Women With ADHD: Evidence From Claims Data. J Clin Psychiatry 81, (2020).

57. A. Gough, J. Morrison, Managing the comorbidity of schizophrenia and ADHD. J Psychiatry Neurosci 41, 150251 (2016).

58. A. A. Grace, Dysregulation of the dopamine system in the pathophysiology of schizophrenia and depression. Nat Rev Neurosci 17, 524–532 (2016).

59. D. A. Martin, D. Marona-Lewicka, D. E. Nichols, C. D. Nichols, Chronic LSD alters gene expression profiles in the mPFC relevant to schizophrenia. Neuropharmacology 83, 1–8 (2014).

60. S. Liang, Q. Wang, X. Kong, W. Deng, X. Yang, X. Li, Z. Zhang, J. Zhang, C. Zhang, X. M. Li, X. Ma, J. Shao, A. J. Greenshaw, T. Li, White Matter Abnormalities in Major Depression Biotypes Identified by Diffusion Tensor Imaging. Neurosci Bull 35, 867–876 (2019).

61. I. Maze, H. E. Covington, 3rd, D. M. Dietz, Q. LaPlant, W. Renthal, S. J. Russo, M. Mechanic, E. Mouzon, R. L. Neve, S. J. Haggarty, Y. Ren, S. C. Sampath, Y. L. Hurd, P. Greengard, A. Tarakhovsky, A. Schaefer, E. J. Nestler, Essential role of the histone methyltransferase G9a in cocaine-induced plasticity. Science 327, 213–216 (2010).

62. A. Sanchez-Gonzalez, E. Thougaard, C. Tapias-Espinosa, T. Canete, D. Sampedro-Viana, J. M. Saunders, R. Toneatti, A. Tobena, J. Gonzalez-Maeso, S. Aznar, A. Fernandez-Teruel, Increased thin-spine density in frontal cortex pyramidal neurons in a genetic rat model of schizophrenia-relevant features. Eur Neuropsychopharmacol (2021).

63. J. F. Lopez-Gimenez, M. T. Vilaro, J. M. Palacios, G. Mengod, Mapping of 5-HT2A receptors and their mRNA in monkey brain: [3H]MDL100,907 autoradiography and in situ hybridization studies. J Comp Neurol 429, 571–589 (2001).

64. R. L. Jakab, P. S. Goldman-Rakic, 5-Hydroxytryptamine2A serotonin receptors in the primate cerebral cortex: possible site of action of hallucinogenic and antipsychotic drugs in pyramidal cell apical dendrites. Proc Natl Acad Sci U S A 95, 735–740 (1998).

65. L. P. Cameron, R. J. Tombari, J. Lu, A. J. Pell, Z. Q. Hurley, Y. Ehinger, M. V. Vargas, M. N. McCarroll, J. C. Taylor, D. Myers-Turnbull, T. Liu, B. Yaghoobi, L. J. Laskowski, E. I. Anderson, G. Zhang, J. Viswanathan, B. M. Brown, M. Tjia, L. E. Dunlap, Z. T. Rabow, O. Fiehn, H. Wulff, J. D. McCorvy, P. J. Lein, D. Kokel, D. Ron, J. Peters, Y. Zuo, D. E. Olson, A non-hallucinogenic psychedelic analogue with therapeutic potential. Nature 589, 474–479 (2021).

66. K. S. Hideshima, A. Hojati, J. M. Saunders, D. M. On, M. de la Fuente Revenga, J. M. Shin, A. Sanchez-Gonzalez, C. M. Dunn, A. B. Pais, A. C. Pais, M. F. Miles, J. T. Wolstenholme, J. Gonzalez-Maeso, Role of mGlu2 in the 5-HT2A receptor-dependent antipsychotic activity of clozapine in mice. Psychopharmacology (Berl) (2018).

67. A. B. Pais, A. C. Pais, G. Elmisurati, S. H. Park, M. F. Miles, J. T. Wolstenholme, A Novel Neighbor Housing Environment Enhances Social Interaction and Rescues Cognitive Deficits from Social Isolation in Adolescence. Brain Sci 9, (2019).

68. P. Zanos, J. N. Highland, B. W. Stewart, P. Georgiou, C. E. Jenne, J. Lovett, P. J. Morris, C. J. Thomas, R. Moaddel, C. A. Zarate, Jr., T. D. Gould, (2R,6R)-hydroxynorketamine exerts mGlu2 receptor-dependent antidepressant actions. Proc Natl Acad Sci U S A 116, 6441–6450 (2019).

69. M. de la Fuente Revenga, D. Ibi, J. M. Saunders, T. Cuddy, M. K. Ijaz, R. Toneatti, M. Kurita, T. Holloway, L. Shen, J. Seto, M. G. Dozmorov, J. Gonzalez-Maeso, HDAC2-dependent Antipsychotic-like Effects of Chronic Treatment with the HDAC Inhibitor SAHA in Mice. Neuroscience 388, 102–117 (2018).

70. M. Kurita, T. Holloway, A. Garcia-Bea, A. Kozlenkov, A. K. Friedman, J. L. Moreno, M. Heshmati, S. A. Golden, P. J. Kennedy, N. Takahashi, D. M. Dietz, G. Mocci, A. M. Gabilondo, J. Hanks, A. Umali, L. F. Callado, A. L. Gallitano, R. L. Neve, L. Shen, J. D. Buxbaum, M. H. Han, E. J. Nestler, J. J. Meana, S. J. Russo, J. Gonzalez-Maeso, HDAC2 regulates atypical antipsychotic responses through the modulation of mGlu2 promoter activity. Nat Neurosci 15, 1245–1254 (2012).

71. B. B. Lake, R. Ai, G. E. Kaeser, N. S. Salathia, Y. C. Yung, R. Liu, A. Wildberg, D. Gao, H. L. Fung, S. Chen, R. Vijayaraghavan, J. Wong, A. Chen, X. Sheng, F. Kaper, R. Shen, M. Ronaghi, J. B. Fan, W. Wang, J. Chun, K. Zhang, Neuronal subtypes and diversity revealed by single-nucleus RNA sequencing of the human brain. Science 352, 1586–1590 (2016).

72. B. Langmead, C. Trapnell, M. Pop, S. L. Salzberg, Ultrafast and memory-efficient alignment of short DNA sequences to the human genome. Genome Biol 10, R25 (2009).

73. Y. Zhang, T. Liu, C. A. Meyer, J. Eeckhoute, D. S. Johnson, B. E. Bernstein, C. Nusbaum, R. M. Myers, M. Brown, W. Li, X. S. Liu, Model-based analysis of ChIP-Seq (MACS). Genome Biol 9, R137 (2008).

74. C. S. Ross-Innes, R. Stark, A. E. Teschendorff, K. A. Holmes, H. R. Ali, M. J. Dunning, G. D. Brown, O. Gojis, I. O. Ellis, A. R. Green, S. Ali, S. F. Chin, C. Palmieri, C. Caldas, J. S. Carroll, Differential oestrogen receptor binding is associated with clinical outcome in breast cancer. Nature 481, 389–393 (2012).

75. M. I. Love, W. Huber, S. Anders, Moderated estimation of fold change and dispersion for RNA-seq data with DESeq2. Genome Biol 15, 550 (2014).

76. Y. Benjamini, Y. Hochberg, Controlling the False Discovery Rate: A Practical and Powerful Approach to Multiple Testing. Journal of the Royal Statistical Society. Series B (Methodological) 57, 289–300 (1995).

77. P. J. Rousseeuw, Silhouettes: A graphical aid to the interpretation and validation of cluster analysis. Journal of Computational and Applied Mathematics 20, 53–65 (1987).

78. D. U. Gorkin, I. Barozzi, Y. Zhao, Y. Zhang, H. Huang, A. Y. Lee, B. Li, J. Chiou, A. Wildberg, B. Ding, B. Zhang, M. Wang, J. S. Strattan, J. M. Davidson, Y. Qiu, V. Afzal, J. A. Akiyama, I. Plajzer-Frick, C. S. Novak, M. Kato, T. H. Garvin, Q. T. Pham, A. N. Harrington, B. J. Mannion, E. A. Lee, Y. Fukuda-Yuzawa, Y. He, S. Preissl, S. Chee, J. Y. Han, B. A. Williams, D. Trout, H. Amrhein, H. Yang, J. M. Cherry, W. Wang, K. Gaulton, J. R. Ecker, Y. Shen, D. E. Dickel, A. Visel, L. A. Pennacchio, B. Ren, An atlas of dynamic chromatin landscapes in mouse fetal development. Nature 583, 744–751 (2020).

79. G. Yu, L. G. Wang, Q. Y. He, ChIPseeker: an R/Bioconductor package for ChIP peak annotation, comparison and visualization. Bioinformatics 31, 2382–2383 (2015).

80. G. Yu, L. G. Wang, Y. Han, Q. Y. He, clusterProfiler: an R package for comparing biological themes among gene clusters. OMICS 16, 284–287 (2012).

81. S. Heinz, C. Benner, N. Spann, E. Bertolino, Y. C. Lin, P. Laslo, J. X. Cheng, C. Murre, H. Singh, C. K. Glass, Simple combinations of lineage-determining transcription factors prime cis-regulatory elements required for macrophage and B cell identities. Mol Cell 38, 576–589 (2010).

82. D. Kim, B. Langmead, S. L. Salzberg, HISAT: a fast spliced aligner with low memory requirements. Nat Methods 12, 357–360 (2015).

83. Y. Liao, G. K. Smyth, W. Shi, featureCounts: an efficient general purpose program for assigning sequence reads to genomic features. Bioinformatics 30, 923–930 (2014).

84. B. Zhang, S. Horvath, A general framework for weighted gene co-expression network analysis. Stat Appl Genet Mol Biol 4, Article17 (2005).

85. P. Shannon, A. Markiel, O. Ozier, N. S. Baliga, J. T. Wang, D. Ramage, N. Amin, B. Schwikowski, T. Ideker, Cytoscape: a software environment for integrated models of biomolecular interaction networks. Genome Res 13, 2498–2504 (2003).

