## Supplementary Figures for "Prolonged epigenetic and synaptic plasticity alterations following single exposure to a psychedelic in mice"

**Supplementary figures and tables**

**fig. S1. Contextual fear extinction in *5-HT$_{2A}$R$^{+/+}$* and *5-HT$_{2A}$R$^{-/-}$* mice.** Fear conditioning (n = 11-26 mice per group; conditioning, F[6,252] = 118.1, p < 0.001; genotype, F[1,252] = 18.28, p < 0.001), expression (n = 11-26 mice per group, $t_{36}$ = 1.46, p > 0.05), generalization (n = 6-15 mice per group, $t_{19}$ = 0.77, p > 0.05), extinction (n = 6-15 mice per group; extinction, F[3,76] = 10.34, p < 0.001; genotype, F[1,76] = 18.69, p < 0.001). Statistical analysis was performed using two-way ANOVA with Sidak's multiple comparison test (conditioning and extinction) or Student's *t* test (expression and generalization). *p < 0.05, **p < 0.01, ***p < 0.001, n.s., not significant. Data correspond to those presented in Figs. 1i,j.

**fig. S2. Experimental set up of treatment (i.p.) with DOI (2 mg/kg), or vehicle.** See also Fig. 3

**fig. S3.** Selected genes (*Drd1* and *Kpna3*) with high correlation between enhancer activity and gene expression level.

**fig. S4. Gene co-expression modules (green, magenta, and turquoise) associated with administration of DOI**
**(A,D,G)** Heatmap of normalized gene expression profiles in the co-expression module (top). The module eigengene values across samples in four experimental groups (bottom).
**(B,E,H)** Selected top categories from GO enrichment analysis.
**(C,F,I)** Visualization of the intramodular connections among the top 100 hub genes in each module. The top 5 genes are in large size and colored orange. The genes involved in the top 15 GO terms are labeled.
(**J**) Each row corresponds to the module eigengene, column to the status of DOI administration. The two numbers in each cell represented the corresponding correlation coefficient and pvalue.

**Table S1.** Metadata on ChIP-seq and mRNA-seq data sets.

**Table S2.** Enriched gene ontology terms associated with the enhancer clusters.

**Table S3.** Enriched transcription factor binding motifs associated with the enhancer clusters.

**Table S4.** A list of genes with high (> 0.4) Spearman correlation between enhancer intensity and expression level.

**Table S5.** Depression SNPs that overlap with differential H3K27ac peaks.

**Table S6.** The eigengene value of each module.

**Table S7.** The list of top 100 hub genes in each module.

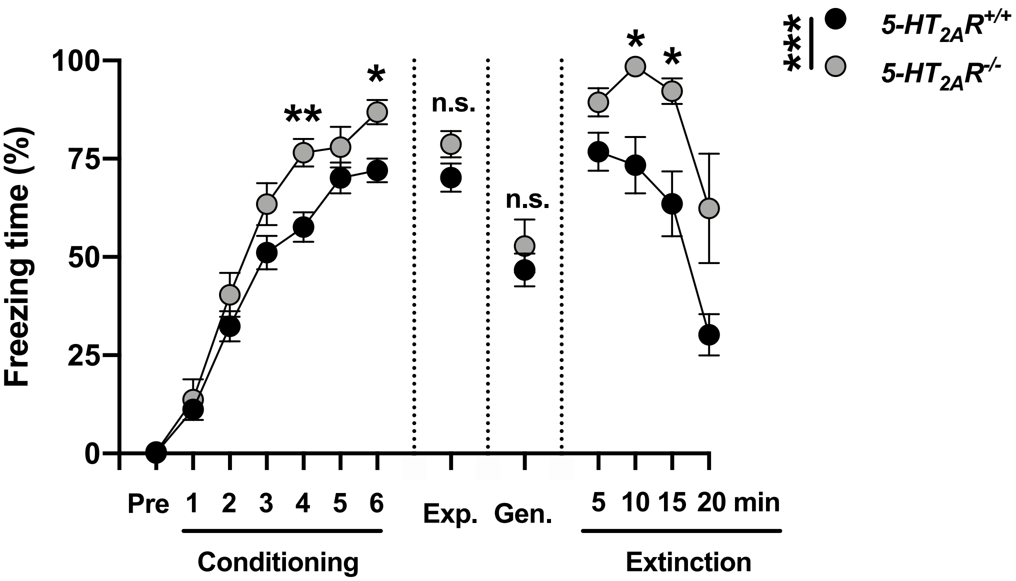


de la Fuente et al (Supplementary Fig 1)

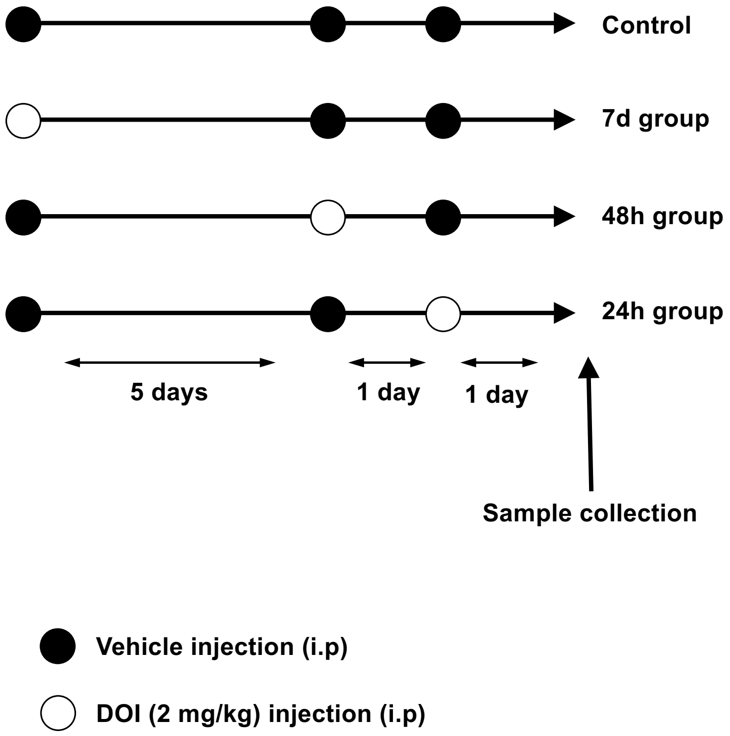


de la Fuente et al (Supplementary Fig 2)

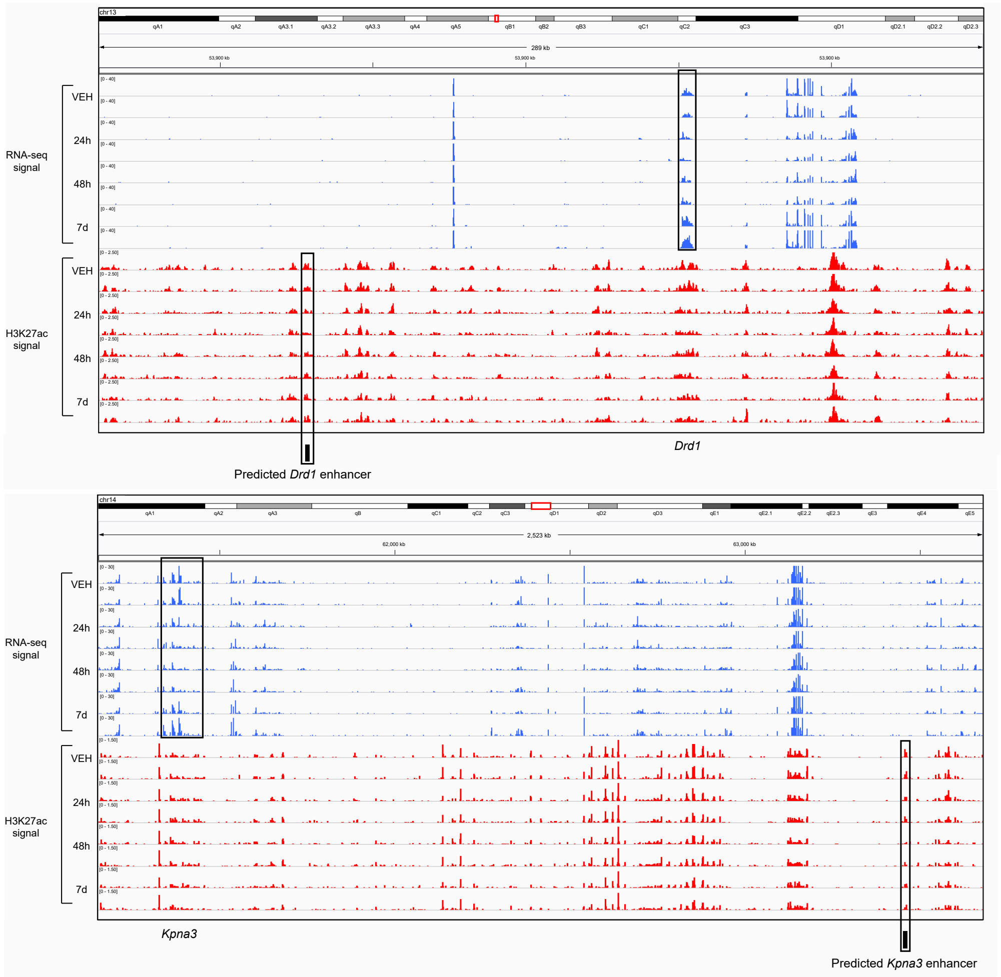


**de la Fuente et al (Supplementary Fig 3)**

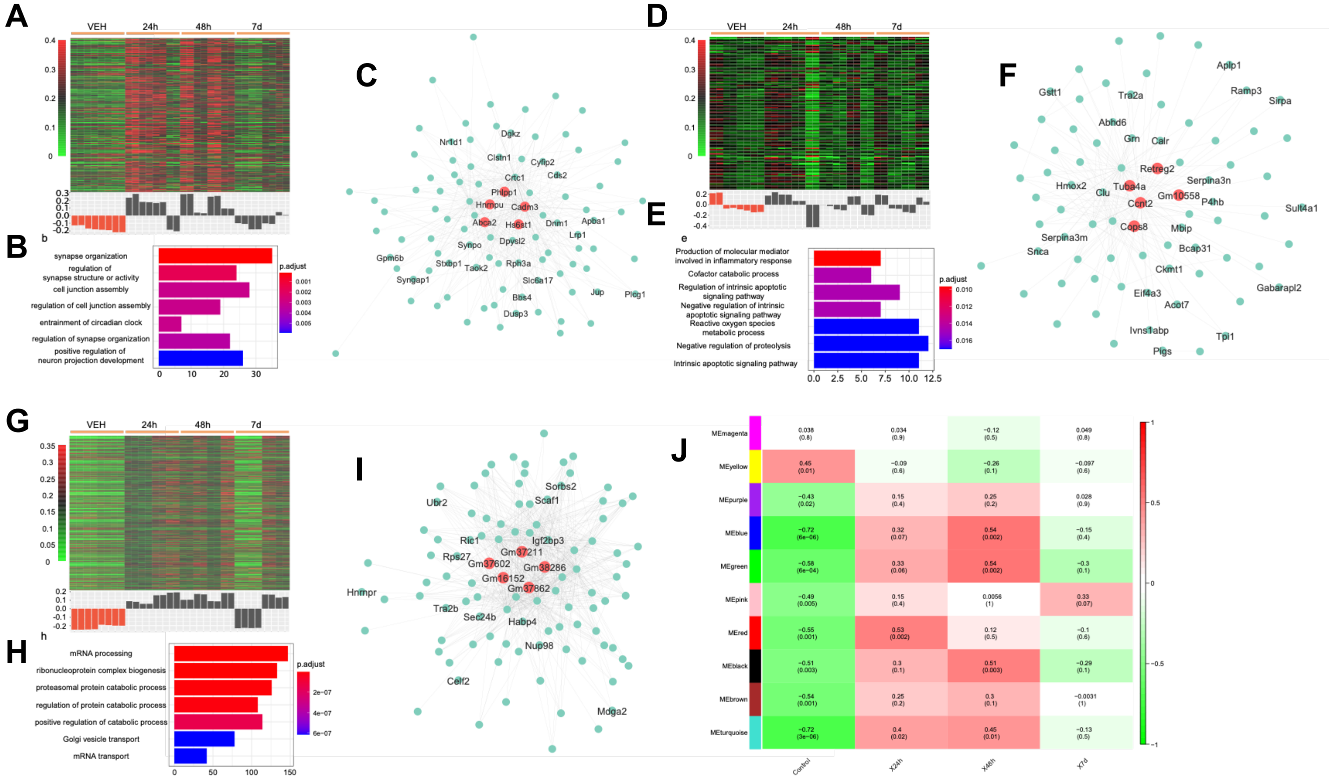


de la Fuente et al (Supplementary Fig 4)
