## Supplementary Table 5 for "Prolonged epigenetic and synaptic plasticity alterations following single exposure to a psychedelic in mice"

| H3K27ac Peaks | | | SNP |  | SNP-linked Gene(s) | |  | Peak-linked Gene(s)^a^ | |
| --- | --- | --- | --- | --- | --- | --- | --- | --- | --- |
| Chr | Start | End |  |  |  |  |  |  |  |
| chr4 | 150581633 | 150585310 | rs301795 |  | Rere |  |  | Cenps |  |
| chr8 | 54856318 | 54857903 | rs6818081 |  | Gpm6a |  |  | Wdr17 | Gpm6a |
| chr9 | 56701937 | 56704480 | rs7175083 |  | Lingo1 |  |  | Lingo1 | Odf3l1 |
| chr9 | 108205689 | 108207054 | rs4625 |  | Dag1 |  |  | Ccdc36 |  |
| chr9 | 108342157 | 108343246 | rs6774721 |  | Gpx1 | Usp4 |  | Ifrd2 |  |
| chr9 | 111407681 | 111409488 | rs9834970 |  | Linc02033 | Hspd1p6 |  | Mlh1 |  |
| chr11 | 3737382 | 3739575 | rs5997786 |  | Osbp2 |  |  | Drg1 |  |
| chr11 | 26961101 | 26961736 | rs11682175 |  | Actg1p22 |  |  | Vrk2 |  |
| chr11 | 29717791 | 29722462 | rs3806572 |  | Rtn4 |  |  | Rtn4 | 4931440F15Rik |
| chr13 | 83731578 | 83733532 | rs6882046 |  | Linc00461 |  |  | Tmem161b | Mef2c |
| chr14 | 13770834 | 13771932 | rs12108177 |  | Sntn | Ac104162.1 |  | Thoc7 |  |
| chr15 | 71842945 | 71844797 | rs77044609 |  | Col22a1 |  |  | Fam135b | Col22a1 |
| chr19 | 20392919 | 20394522 | rs3758354 |  | Anxa1 | Aldh1a1 |  | Anxa1 |  |
| chr19 | 46907975 | 46910493 | rs10883832 |  | Nt5c2 |  |  | Cyp17a1 | Cnnm2 |

^a^H3K27ac peaks were linked to genes either through correlation with gene expression or to the nearest genes
